# IRF4 is a master regulator of multinucleated giant cell formation

**DOI:** 10.1101/2025.11.03.686191

**Authors:** Melanie Hofmann, Gerwin Heller, Markus Kieler, Laszlo Musiejovsky, Lukas Kremp, Martina Kerndl, Paul Ettel, Birgit Niederreiter, Lucia Quemada Garrido, Anahita Sedighi, Thomas Krausgruber, Leonhard X. Heinz, Axel Dietschmann, David Voehringer, Bart Everts, Thomas Weichhart, Omar Sharif, Stephan Blüml, Gernot Schabbauer

**Affiliations:** Institute of Vascular Biology and Thrombosis Research, Center for Physiology and Pharmacology, Medical University of Vienna, Vienna, Austria; Christian Doppler Laboratory for Arginine Metabolism in Rheumatoid Arthritis and Multiple Sclerosis, Vienna, Austria; Department of Medicine I, Division of Oncology, Medical University of Vienna, Vienna, Austria; Department of Internal Medicine III, Division of Rheumatology, Medical University of Vienna, Vienna, Austria; Center for Pathobiochemistry and Genetics, Medical University of Vienna, Vienna, Austria; Medical University of Vienna, Center for Medical Data Science, Institute of Artificial Intelligence, Vienna, Austria; CeMM Research Center for Molecular Medicine of the Austrian Academy of Sciences, Vienna, Austria; Department of Infection Biology, University Hospital Erlangen and Friedrich-Alexander-Universität Erlangen-Nürnberg, Erlangen, Germany; Junior Research Group Adaptive Pathogenicity Strategies, Leibniz Institute for Natural Product Research and Infection Biology - Hans Knöll Institute, Jena, Germany; Centre for Infectious Diseases, Leiden University Medical Centre, Leiden, Netherlands; Christian Doppler Laboratory for Immunometabolism and Systems Biology of Obesity-Related Diseases (InSpiReD), Vienna, Austria

## Abstract

Multinucleated giant cells (MGCs) are a hallmark pathological feature of a wide range of diseases, yet the mechanisms underlying their formation and function remain poorly understood. Although it is recognized that MGCs arise from a heterogeneous pool of myeloid precursors, how a committed MGC fate develops therein remains unknown.

Here, combining temporal *in vitro* and *in vivo* differentiation of bone marrow-derived myeloid precursors with single cell and bulk RNA sequencing as well as CRISPR/Cas9-mediated gene editing, we shed insight into how MGCs emerge. Our findings reveal that coordinated upregulation of cell fusion genes and cellular metabolism, particularly oxidative phosphorylation, regulate the transition from macrophage progenitors to fusion competent cells. *In vivo* fate-mapping unveils *Ms4a3*-, and *Cd11c*-but not *Cx3cr1*-traced cells as the predominant precursor populations for MGCs in a lung granuloma formation model. Notably, transcription factor profiling in the progression from early myeloid precursors to pre-MGCs identifies IRF4 as a key molecular switch driving MGC generation. IRF4^+^ MGCs are present in different pathologies, including *Schistosoma mansoni* egg induced granulomas, *Aspergillus fumigatus* conidia mediated allergic airway inflammation and human head and neck squamous cell carcinomas. Mechanistically, IRF4 controls critical fusion related genes such as *Dcstamp* and *Ocstamp*. Consequently, *Irf4* deficient cells are unable to develop into MGCs.

Collectively, our work delineates the trajectory of MGC differentiation, establishing IRF4 as a defining transcription factor required for the generation of fusion-competent progenitors that ultimately give rise to MGCs.

## Introduction

Multinucleated giant cells (MGCs) are striking morphological entities found in various pathological contexts and are characterized by their large size and the presence of multiple nuclei ^1, 2, 3^. While their appearance is often indicative of chronic inflammation or specific disease states, the mechanisms of their formation as well as their exact functions are not well understood and therefore subjects of ongoing scientific investigation ^2, 4, 5^. MGCs can be subdivided into distinct forms, the most common types including: Foreign body giant cells, which form around non-degradable material, such as sutures, medical implants, inhaled particulate matter but also amyloid deposits; Langhans giant cells that represent hallmarks of granulomatous inflammation such as tuberculosis and sarcoidosis, where they contribute to the containment and modulation of the immune response or wall off persistent infectious agents like *Mycobacterium tuberculosis*, *Schistosoma mansoni* (*S. mansoni*) and others ^6, 7, 8, 9, 10^ and osteoclasts, which are centrally involved in bone homeostasis ^11^. The formation of MGCs primarily occurs through cell fusion where individual myeloid cells, stimulated by specific cytokines (e.g., interleukin (IL)-4, IL-13, interferon (IFN)-γ, tumor necrosis factor (TNF)-α, receptor activator of nuclear factor kappa-Β ligand (RANKL)) or in the presence of persistent foreign material, undergo membrane fusion ^12, 13^. This complex process involves the upregulation of adhesion molecules and specific fusogenic proteins, most prominently dendrocyte expressed seven transmembrane protein (DC-STAMP) and osteoclast stimulatory transmembrane protein (OC-STAMP) ^14, 15^. In some cases, particularly observed in tuberculosis associated MGCs, a single cell may undergo repeated nuclear division (karyokinesis) without subsequent cytoplasmic separation (cytokinesis) ^16^.

Osteoclasts (OCs), vital for maintaining bone homeostasis by resorbing aged or damaged bone tissue and therefore crucial for bone growth, repair, and mineral balance, remain the best studied type of MGCs. Major factors regulating their differentiation such as the RANK/RANKL/(Osteoprotegerin) OPG system, costimulatory molecules and transcription factors conferring osteoclast identity (most prominently nuclear factor of activated T-cells 1 (NFATc1)) as well as many aspects of their cellular origin have been elucidated over the last decades ^11, 17, 18, 19, 20, 21^. In contrast, little is known about the molecular and cellular mechanisms that lead to IL-4 induced MGCs, apart from the fact that they are of myeloid origin and, akin to OCs, require the downregulation of macrophage-associated gene signatures as well as the upregulation of essential fusogens like DC-STAMP and OC-STAMP as prerequisites for their formation^8, 13, 14, 22, 23, 24^. Therefore, a deeper understanding of the generation and function of these MGC type is essential and will further aid in elucidating their contribution to various diseases.

## Results

### Transcriptional and metabolic profiling of MGC differentiation

To characterize the prerequisites for multinucleated giant cell (MGC) formation in more detail, we performed time-resolved characterization of an established MGC *in vitro* culture system ^12, 25^. This assay involves the isolation of murine bone marrow (BM) cells and culturing them in the presence of macrophage colony-stimulating factor (M-CSF) for 3 days to enrich for the myeloid precursor cell compartment before inducing MGC development by stimulation with granulocyte-macrophage colony-stimulating factor (GM-CSF) and interleukin (IL)-4 (**Figure 1a**). Using live-cell time-lapse microscopy of a mixture of GFP-(BM cells isolated from UBI-GFP mouse) and tdTomato-expressing cells (BM cells derived from Ai14^fl^ *Ms4a3*-cre mouse), we detected numerous GFP/tdTomato double positive MGCs, confirming that MGC formation in our system requires cell to cell fusion as opposed to failed cytokinesis, which is consistent with previous reports using thioglycolate elicited myeloid cells as precursor population ^25, 26^. These time-lapse series further revealed that fusion starts about 12 hours post stimulation with GM-CSF and IL-4 and that the fusion rate slows down shortly after 24 hours until the end of the assay at 48 hours (**Figure 1b** and **Suppl. Figure 1a and b**). This cell fusion dynamics suggests that most myeloid precursor cells have completed the differentiation towards a fusogenic cell state between 12-24 hours. To identify fusion-mediating gene signatures in a time-dependent manner, we initially performed a principal component analysis (PCA) of bulk RNA sequencing (bulk RNA-seq) of myeloid precursor cells at different stages post fusion induction (**Figure 1c**). Focusing on time points co-inciding with cellular fusion (12h and 24h), we observed substantial transcriptional alterations (ca. 6000 deregulated genes) in comparison to the precursor state (0h). All shared differentially regulated genes (3179) between the 0h vs 12h and 0h vs 24h conditions were considered as putative fusion genes, given within these two timepoints most cells are in a fusogenic/fusion-competent cell state (**Figure 1d**). Kinetic clustering therein revealed an early and a late upregulated gene cluster comprising 596 and 980 genes, respectively (**Figure 1e**), as well as an early and a late downregulated cluster including 733 and 888 genes, respectively (**Suppl. Figure 1c**). In line with previously published data ^22^, MGC differentiation was characterized by downregulation of macrophage-lineage associated genes like MAF bZIP transcription factor B (*Mafb*), interferon regulatory factor (*Irf*)*8* and the colony stimulating factor 1 receptor (*Csf1r*) (**Suppl. Figure 1c and d**). GO term enrichment analysis identified several processes related to response to (immune) stimuli, including activation of the JAK-STAT pathway as a consequence of the GM-CSF and IL-4 stimulation in the early upregulated gene cluster (**Figure 1e**, upper panel). Cadherin 1 (*Cdh1*), that has been shown to be important for MGC formation ^27^, was among the early induced genes, as well as other MGC associated markers like matrix metallopeptidase (*Mmp*)*13* and arginase 1 (*Arg1*), which are less well established ^28^. In contrast, prominent fusion factors such as *Dcstamp* and *Ocstamp*, were among the genes of the late upregulated cluster (**Figure 1e**, lower panel) ^14, 15^. GO term enrichment analysis of these later induced genes was dominated by processes related to phagocytosis, cell migration, adenosine triphosphate (ATP) biosynthesis, mitochondrial respiration and oxidative phosphorylation (OXPHOS), suggesting increased energy demands of myeloid precursor cells during differentiation towards MGCs (**Figure 1e**, lower panel). Therefore, we set out to analyze the metabolic properties of these differentiating cells by measuring oxygen consumption rate (OCR) as well as the extracellular acidification rate (ECAR) at different timepoints post fusion induction (**Figure 1f**). We found a continuous increase in both OCR and ECAR in the course of MGC formation (**Figure 1f** and **Suppl. Figure 1e**), in line with an increase in total ATP production rate (**Figure 1g**). However, the proportion of mitochondrial ATP production (**Figure 1h**) as well as the basal OCR levels (**Suppl. Figure 1e**) were significantly enhanced during MGC fate commitment, suggesting that pre-MGCs favor mitochondrial respiration as their main energy generation pathway. To assess whether glycolysis or oxidative phosphorylation are functionally required for the differentiation of MGCs, we induced fusion in the presence of different mitochondrial complex inhibitors like oligomycin, antimycin A, rotenone or 2-Thenoyltrifluoroacetone (TTFA) and the glycolysis blocking agent 2-Deoxy-D-glucose (2-DG). We found that inhibitors of both pathways of ATP generation, mitochondrial respiration as well as glycolysis, impaired MGC development, with mitochondrial inhibitors having a more pronounced blocking effect compared to 2-DG or glucose depletion (**Figure 1i and j, Suppl. Figure 1f and g**). Taken together, these results suggest that an increase in cellular metabolism, particularly OXPHOS, is tightly linked to successful MGC generation.

**Figure 1:**
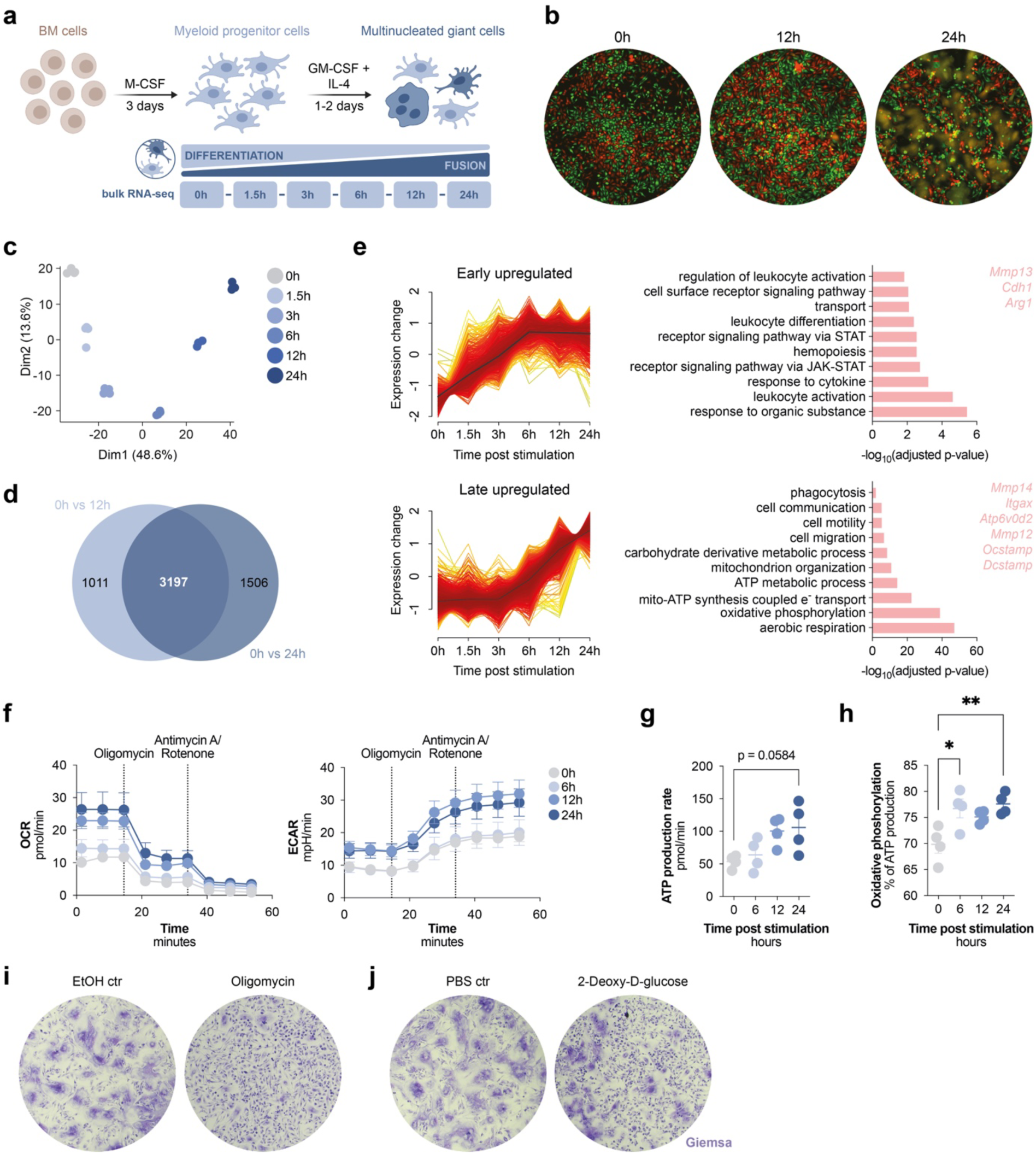
Transcriptional and metabolic profiling of MGC differentiation. **(a)** Schematic overview of the *in vitro* formation assay of MGCs derived from murine BM cells including the depiction of timepoints for bulk RNA-seq sample generation. **(b)** Representative images of live-cell time-lapse microscopy of a mixture of UBI-GFP (green) and Ai14^fl^ *Ms4a3*-cre (red) BM cells stimulated with IL-4 and GM-CSF for 0h, 12h and 24h. Fused cells are depicted in yellow (overlap of red and green signal). **(c)** Principal component analysis (PCA) plot of all bulk RNA-seq samples. **(d)** Venn diagram depicting the overlap proportion of differentially expressed genes (DEGs) between 0h vs 12h and 0h vs 24h post fusion induction (putative fusion genes). **(e)** Unbiased gene expression pattern analysis plots of the 3197 putative fusion genes shown in **(d)** as well as significantly enriched pathways described by genes included in the respective clusters as determined by geneset over-representation analysis (ORA) based on the GO biological pathway (GO:BP) database. Upper and lower panel represent the early and late upregulated gene trajectories and their respective selections of significantly enriched GO:BP pathways as well as a selection of genes included in these trajectories, respectively. **(f)** Oxygen consumption rate (OCR) and extracellular acidification rate (ECAR) of MGC precursor cells at 0h, 6h, 12h and 24h post fusion induction. **(g)** and **(h)** Total ATP production rate as well as the proportion of ATP production generated by oxidative phosphorylation of myeloid precursor cells stimulated with IL-4 and GM-CSF for the indicated timepoints. **(i)** and **(j)** Representative images of the inhibiting effect of either 50nM Oligomycin **(i)** or 200µM 2-DG **(j)** on MGC formation of murine BM cells including solvent controls. **(b)** Representative images of in total n=2 biological replicates of 2 independent experiments. **(c-e)** Data represent n=4 biological replicates per timepoint. **(f-h)** Data represent n=4 biological replicates per timepoint of 2 independent experiments. **(i)** and **(j)** Representative images of in total n=4 biological replicates of 2 independent experiments. Data are mean ± SEM, *p < 0.05, **p < 0.01, one-way ANOVA post-hoc pairwise comparisons with Tukey correction.

### Single cell characterization of fusion competent cells

The above bulk analyses cannot rule out the possibility that some of the changes in transcriptome and metabolism reflect shifts in relative cell composition (early on few MGCs, later many more) rather than cell intrinsic changes in these parameters. To address this, we performed single cell RNA sequencing (scRNA-seq) of BM cells cultured under MGC forming conditions (**Figure 2a**). The time schedule for the preparation of scRNA-seq samples was adjusted and optimized to use mononucleated pre-MGCs, alleviating that cells would be too large and burst during harvest, but already display significant upregulation of known fusion associated genes, which occurs between the 0h and 12h timepoint (**Figure 1e**). Uniform manifold approximation and projection (UMAP) visualization of the integrated transcriptomics data from all three timepoints post fusion induction demonstrated that akin to already published data on the trajectory of osteoclast differentiation ^29^, BM cells cultured in the presence of M-CSF (0h timepoint) displayed a certain degree of cellular heterogeneity, that is composed of proliferating (proliferating) and non-proliferating (precursor) myeloid progenitor cells, a cluster defined as monocyte-dendritic cell like cells (moDC-like) and neutrophils (**Figure 2b and c, Suppl. Figure 2a**). During differentiation, one cluster (referred to as fusion competent (FC)) appeared 6h post fusion induction and the proportion of cells associated with this cluster increased substantially over time, while the percentage of cells composing the precursor cluster decreased (**Figure 2b and c**), suggesting that precursor cells would give rise to fusion competent cells. Genes known to be essential for MGC formation like *Dcstamp*, *Ocstamp* and *Cdh1* were enriched in the fusion competent cluster (**Figure 2d**, upper panel). In contrast, macrophage associated genes such as *Csfr1*, the mannose receptor C-type 1 (*Mrc1*) and *Mafb* were downregulated, as previously described ^22^ (**Figure 2d**, lower panel). KEGG enrichment analysis of differentially expressed genes (DEGs) of the fusion competent vs the myeloid precursor cell cluster demonstrated processes related to adhesion and cytoskeletal organization, chemotaxis and migration, energy generation and metabolism, with a notable upregulation of genes involved in OXPHOS, as well as phagocytosis and efferocytosis as overrepresented pathways (**Figure 2e** and **Suppl. Figure 2b**). Therefore confirming, that cell intrinsic transcriptional changes leading to a fusion competent cell state are occurring prior to myeloid cell fusion.

**Figure 2:**
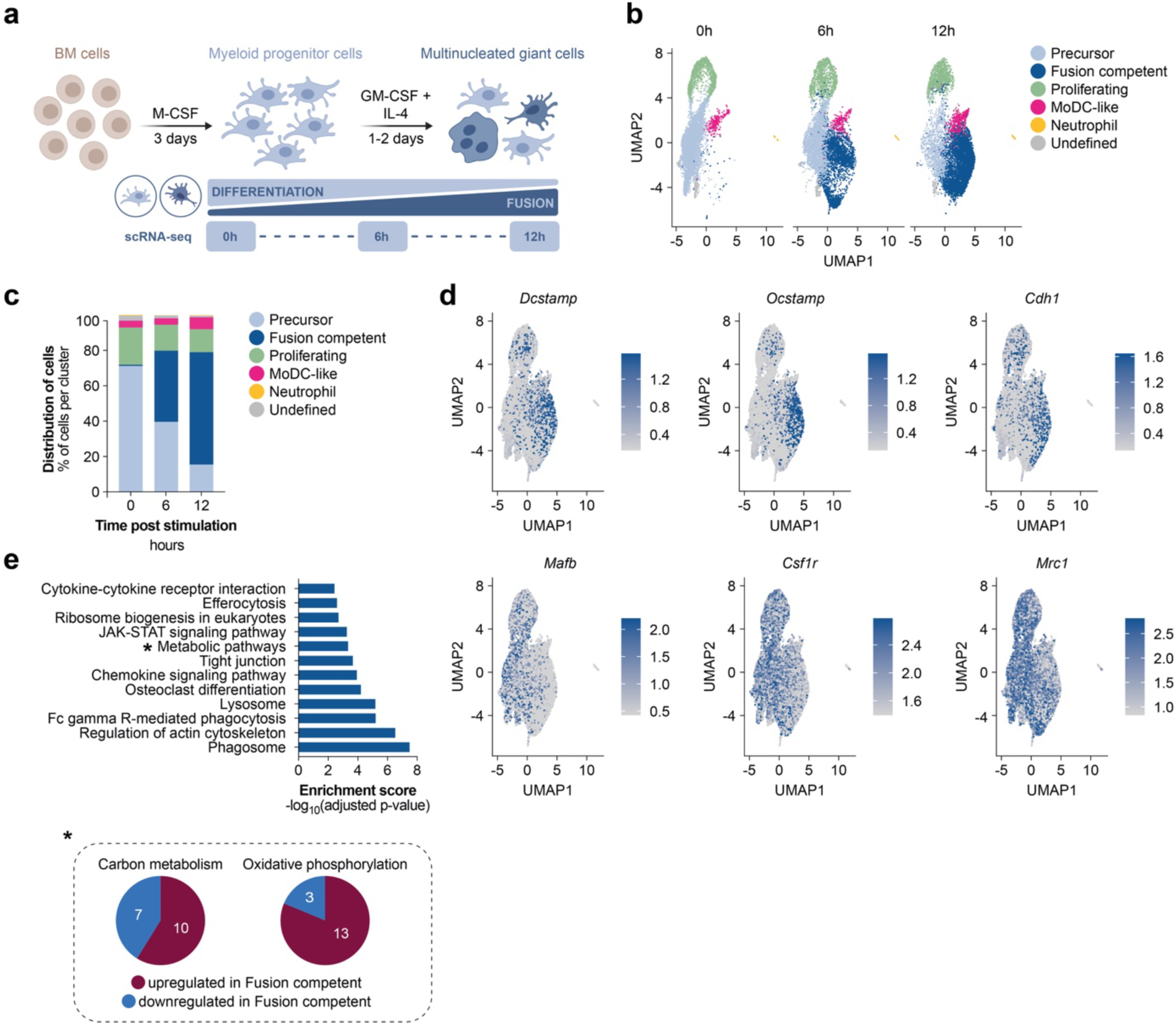
Single cell characterization of fusion competent cells. **(a)** Schematic overview of the *in vitro* MGC formation assay and outline of sample harvest for scRNA-seq data acquisition. **(b)** UMAP plot visualization of clusters that were defined by an unsupervised graph-based clustering approach covering the three different timepoints. **(c)** Bar graph depicting the percentages of cells per cluster and per timepoint. **(d)** The expression pattern of genes essential for MGC formation *Dcstamp*, *Ocstamp* and *Cdh1* (upper panel) and known MGC precursor genes *Mafb*, *Csf1r* and *Mrc1* (lower panel) in the UMAP visualization. **(e)** Selection of significantly enriched pathways of DEGs of precursor vs fusion competent cells as determined by ORA based on the KEGG database. Pie charts additionally depict the number of significantly up- and downregulated genes included in the indicated metabolic pathways (carbon metabolism and oxidative phosphorylation) specific for the fusion competent cluster. **(b-e)** Data represent n=2 pooled biological replicates.

### Origin and function of MGCs *in vivo*

To test whether similar mechanisms and characteristics of MGC formation operate *in vivo* we employed a model of MGC associated lung granuloma development using *S. mansoni* eggs (**Figure 3a**)^30^. Upon *i.v.* administration of *S. mansoni* eggs, cytokines required for MGC generation, IL-4 and GM-CSF, were markedly upregulated in lung tissues, as were the mRNA expression levels of *Dcstamp* (**Figure 3b and c**). Since specific antibodies for known MGC markers like DC-STAMP and OC-STAMP are not available and MGCs are hard to distinguish from large cell infiltrates on H&E-stained histological slides, we first set out to establish a way of robustly detecting MGCs. In both our bulk RNA-seq and scRNA-seq dataset, *Arg1*, encoding for an enzyme previously demonstrated to be associated with MGCs ^28^, was one of the highest induced genes post fusion induction (**Figure 1e**, **Suppl. Figure 1d** and **Suppl. Figure 3a**). Immunofluorescence staining for this enzyme of *in vitro* differentiated MGCs confirmed the presence of ARG1 on a protein level and the usability as an MGC marker (**Suppl. Figure 3b**).

**Figure 3:**
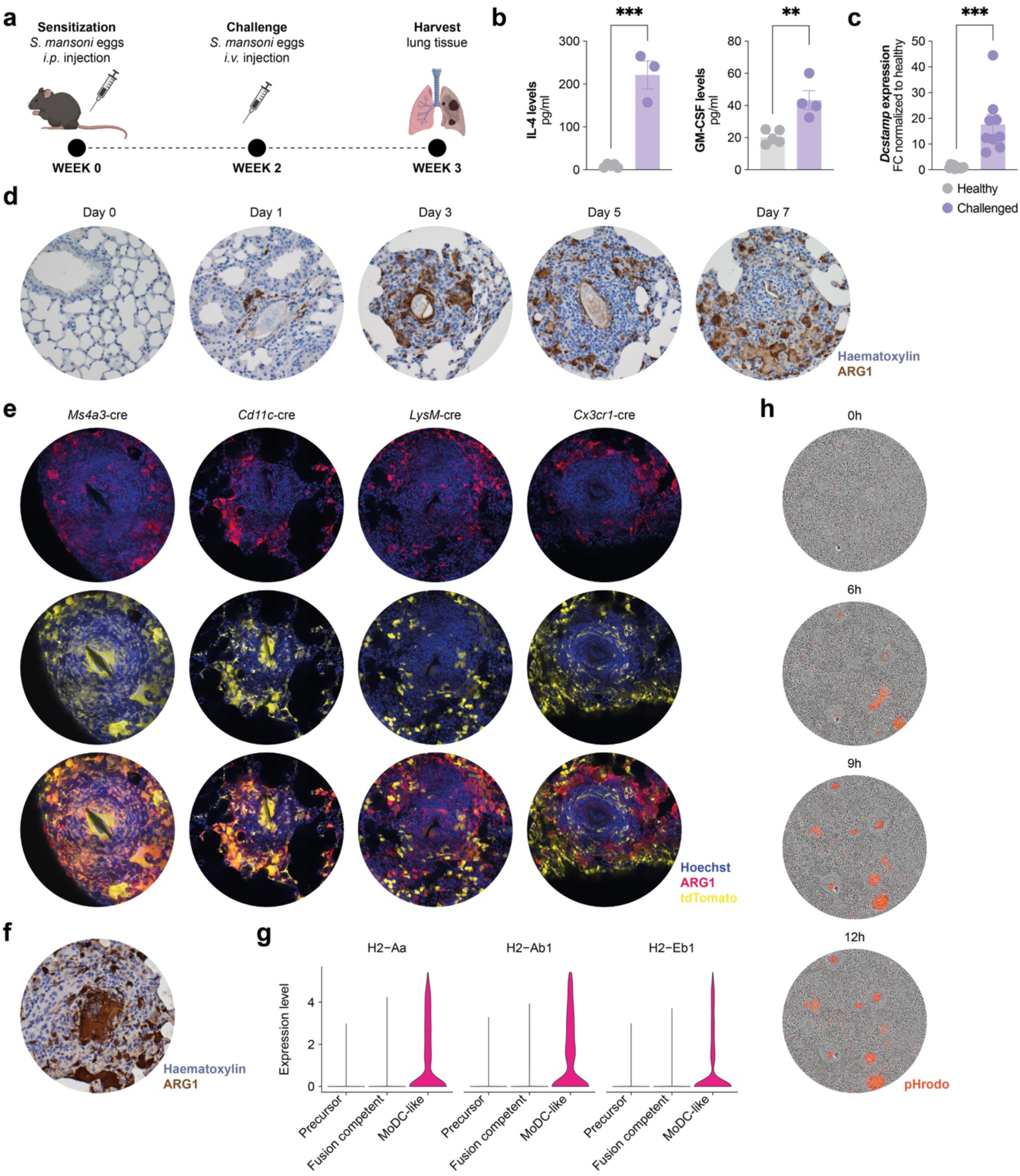
Origin and function of MGCs *in vivo*. **(a)** Schematic overview of *S. mansoni* egg induced granuloma and MGC formation *in vivo*. **(b)** IL-4 and GM-CSF levels in the lung tissue of healthy and *S. mansoni* egg challenged mice on day 7 post *i.v.* injection. **(c)** Relative mRNA expression of *Dcstamp* in lung tissue of healthy and *S. mansoni* egg challenged animals on day 7 post *i.v.* injection. **(d)** Representative pictures of ARG1 staining in lungs on various timepoints post *i.v.* injection with *S. mansoni* eggs. **(e)** Images of lung granulomas (day 7) of different fate mapping mouse strains stained for ARG1 (AF647, red) in upper panel and detection of endogenous tdTomato signal (yellow) in the middle panel with Hoechst counter staining (blue). Lower panel depicts overlay of ARG1 and tdTomato signal. **(f)** Representative image of ARG1 stained lung granuloma sections depicting an MGC containing engulfed granulocytes. **(g)** Violin plots depicting the expression levels of various MHCII associated genes across different clusters of the scRNA-seq dataset. **(h)** Representative images of fully differentiated murine BM cell derived MGCs phagocytosing apoptotic neutrophils labelled with pHrodo at indicated timepoints post addition of target cells at a ratio of 3:1 (apoptotic cells:MGCs). Red signal indicates successful uptake. **(b)** Data represent n=5 healthy and n=3-4 challenged animals. **(c)** Data represent n=11 healthy and n=9 challenged mice of 3 independent experiments. **(d)** Representative images of in total n=5 animals for day0, n=3 for day1, n=7 for day3, n=7 for day5 and n=6 for day7 post *i.v.* challenge with *S. mansoni* eggs of 2 independent experiments. **(e)** Representative images of lungs 7 days post *i.v.* challenge with *S. mansoni* eggs, n=8 challenged of 2 independent experiments for Ai14^fl^ *Cx3cr1-*cre, n=4 challenged for Ai14^fl^ *Cd11c-*cre, n=5 challenged of 2 independent experiments for Ai14^fl^ *LysM-*cre as well as n=3 challenged for Ai14^fl^ *Ms4a3-*cre animals. **(f)** Representative image of in total at least n=6 animals. **(g)** Data represent n=2 pooled biological replicates. **(h)** Representatives of 5-9 images per well of n=2-3 technical replicates and 2 independent experiments. Data are mean ± SEM, **p < 0.01 and ***p < 0.001, unpaired *t*-test.

Using ARG1 expression in combination with a number of nuclei ≥ 3 to characterize MGCs in lung tissue, we could determine that MGC formation started already 3 days post challenge of mice with *S. mansoni* eggs, suggesting that the differentiation process of myeloid precursor cells towards MGCs *in vivo* occurs with similar kinetics as in our *in vitro* culture system (**Figure 3d**). Due to this rapid reprogramming, we were wondering which type of the myeloid cell lineage would acquire fusion competency and give rise to MGCs. Although monocyte precursors have been implicated as MGC progenitor cells in mice suffering from *Mycobacterium tuberculosis* infection ^8^, evidence exists that also tissue resident alveolar macrophages can commit to MGC formation in the presence of *Aspergillus fumigatus* ^30^. To determine the capacity of different myeloid subsets to serve as precursors for MGCs in our *in vivo* model, we induced lung granuloma formation in several fate mapping mouse lines. Using Ai14^fl^ *Ms4a3*-cre mice, we could demonstrate that cells derived from the granulocyte-monocyte progenitor (GMP) population are crucial precursor cells for MGCs as all multinucleated cells within the lung granuloma were uniformly and brightly positive for both, the MGC marker ARG1 and tdTomato (**Figure 3e**, 1^st^ column, **Suppl. Movie 1**). Similarly, MGCs in Ai14^fl^ *Cd11c*-cre animals were positive for tdTomato, whereas MGCs in Ai14^fl^ *LysM*-cre mice exhibited only a weak tdTomato-positive signal (**Figure 3e**, 2^nd^ and 3^rd^ column, **Suppl. Movie 2 and 3**). Interestingly, data derived from Ai14^fl^ *Cx3cr1*-cre fate mapping mice suggests that *Cx3cr1*-traced cells are not participating in MGC formation as demonstrated by virtual absence of tdTomato in ARG1-positive multinucleated cells (**Figure 3e**, 4^th^ column and **Suppl. Movie 4**). Altogether, this data reveals that in our *in vivo* model, monocyte progenitors and/or peripheral classical monocytes are the main precursors, whereas *Cx3cr1*-traced myeloid cells such as non-classical monocytes and alveolar macrophages hardly contribute to MGC formation.

Notably, in accordance with our bulk and scRNA-seq data highlighting an association of fusion competence to phagocytosis, MGCs in the granulomas surrounding the *S. mansoni* egg showed intracellular accumulation of polymorphonuclear granulocytes implying a role for MGCs in the efferocytosis of these cells (**Figure 3f**). Importantly, pre-MGCs completely lacked MHCII expression (**Figure 3g**), suggesting limited ability of these cells to present antigens to T cells which is in line with the notion of an immunologically silent uptake of cellular debris. When we tested this hypothesis *in vitro,* we found that MGCs are indeed capable of engulfing apoptotic neutrophils as demonstrated by an increase of pHrodo signal due to intracellular accumulation of dying cells over time as well as taking up large numbers of aggregated polymorphonuclear cells suggesting a role for MGCs in the clearing of micro-abscesses (**Figure 3h**, **Suppl. Movies 5 and 6**).

### Transcription factor profiling identifies IRF4 in MGC differentiation

To uncover the molecular drivers of MGC formation, we focused on temporal changes in the gene regulatory networks and identified several transcription factors to be significantly upregulated during differentiation towards a fusion competent cell (**Figure 4a**). Integration with scRNA-seq data revealed *Irf4* as one of the most prominently regulated transcription factors during MGC development that was also specifically associated with a fusion committed cell state as compared to precursor cells (**Figure 4b and c**). In addition to mRNA expression *in vitro* (**Figure 4a-c**) and *in vivo* (**Suppl. Figure 4a**), we verified the upregulation of IRF4 on a protein level during MGC differentiation using immunohistochemistry staining of *in vitro* MGC cultures (**Figure 4d**) and *in vivo* in developing MGCs during granuloma formation in lung sections of *S. mansoni* challenged mice (**Figure 4e** and **Suppl. Figure 4b**). Importantly, we could detect a marked signal for IRF4 in ARG1^+^ MGCs associated with lung granulomas, which was absent in control mice (**Figure 4e**). Additionally, IRF4 expressing ARG1^+^ MGCs were present in lung tissue of an *Aspergillus fumigatus* (*Af*) mediated allergic airway inflammation model, suggesting that IRF4 is involved in the formation of MGCs in a range of pathologies (**Figure 4f**). Of note, IRF4 and concomitant DC-STAMP upregulation was also detected in human monocytes differentiated into MGCs *in vitro* using a variety of different stimuli (**Figure 4g and h**, **Suppl. Figure 4c**). In addition, analyzing MGCs in a spatial transcriptomics dataset of samples from head and neck squamous cell carcinoma patients^31^ revealed the existence of *IRF4*^+^ and *DC-STAMP*^+^ MGCs in human disease (**Figure 4i** and **Suppl. Figure 4d**). Taken together, IRF4 expressing MGCs are found in various pathologies in mice and man.

**Figure 4:**
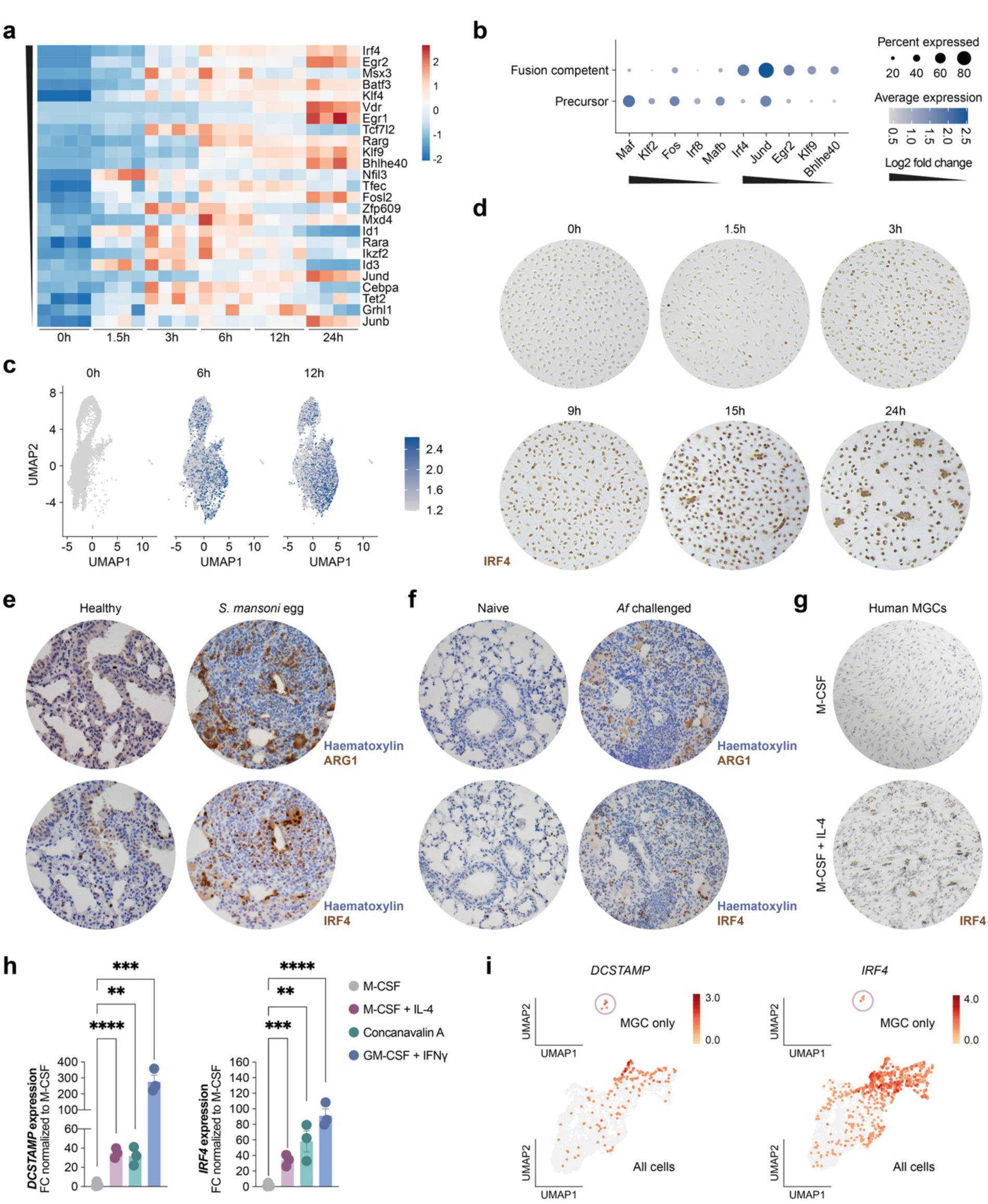
Transcription factor profiling identifies IRF4 in MGC differentiation. **(a)** Heatmap depicting the top 25 significantly upregulated transcription factors of 0h vs 6h samples (after overlaying the DEGs of 0h vs 6h with transcription factor genes included in the transcription factor database TFDB v4.0^50^) at indicated timepoints during the course of MGC formation. Genes are ordered by decreasing shrunk Log2 fold change values, symbolized by black triangle on the left side of the heatmap. **(b)** Dot plot showing the expression levels of the 5 most significantly deregulated transcription factors specific for either the precursor or the fusion competent cell cluster. **(c)** The expression pattern of *Irf4* over the course of development towards fusion competent cells in the UMAP visualization. **(d)** Representative images of IRF4 stained primary murine myeloid precursor cells at various timepoints post fusion induction. **(e)** Representative images of ARG1 and IRF4 stained lung sections of healthy and *S. mansoni* egg challenged mice on day7 of granuloma formation. **(f)** Representative pictures of ARG1 and IRF4 stained lung sections of naive and *Af* challenged animals. **(g)** Representative results of IRF4 staining of CD14^+^ sorted monocytes from human blood stimulated with either M-CSF alone or together with IL-4 to induce MGC formation. **(h)** *DCSTAMP* and *IRF4* mRNA expression levels of human derived monocytes cultured in the presence of M-CSF or various MGC inducing stimuli. **(i)** Expression pattern of *DCSTAMP* and *IRF4* of the MGC^high^ Patient 3 sample derived from a publicly available spatial transcriptomics dataset either depicted for the MGC cluster alone (upper panel) or for all cells together (lower panel) in the UMAP visualization. **(a)** Data represent n=4 biological replicates per timepoint. **(b)** and **(c)** Data represent n=2 pooled biological replicates. **(d)** Representative images of in total n=2 biological replicates. **(e)** Representative data of in total n=7 healthy and n=11 challenged animals of 3 independent experiments. **(f)** Representative images of n=6 mice per group. **(g)** Representative staining of in total 2 blood donors of 2 independent experiments. **(h)** Data represent n=3 technical replicates of one donor as representative of in total 2 individuals. Data are mean ± SEM, **p < 0.01, ***p < 0.001 and ****p < 0.0001 unpaired *t*-test (each condition compared to M-CSF).

### IRF4 specifically regulates cell fusion in MGCs

To functionally analyze the role of IRF4 in MGC development, we generated *Irf4* deficient HoxB8 cells (**Figure 5a**, **Suppl. Figure 5a**). While wildtype HoxB8 cells were able to differentiate into MGCs, upregulate *Dcstamp* and IRF4 expression during the course of MGC formation *in vitro* similar to primary BM cells (**Suppl. Figure 5b and c**), *Irf4* deficient cells were completely incapable of generating multinucleated cells (**Figure 5b**). Expression of IRF4 was even an absolute cell intrinsic requirement for multinucleation, as *Irf4* knockout cells could not fuse with wildtype cells to give rise to MGCs (**Figure 5c**).

**Figure 5:**
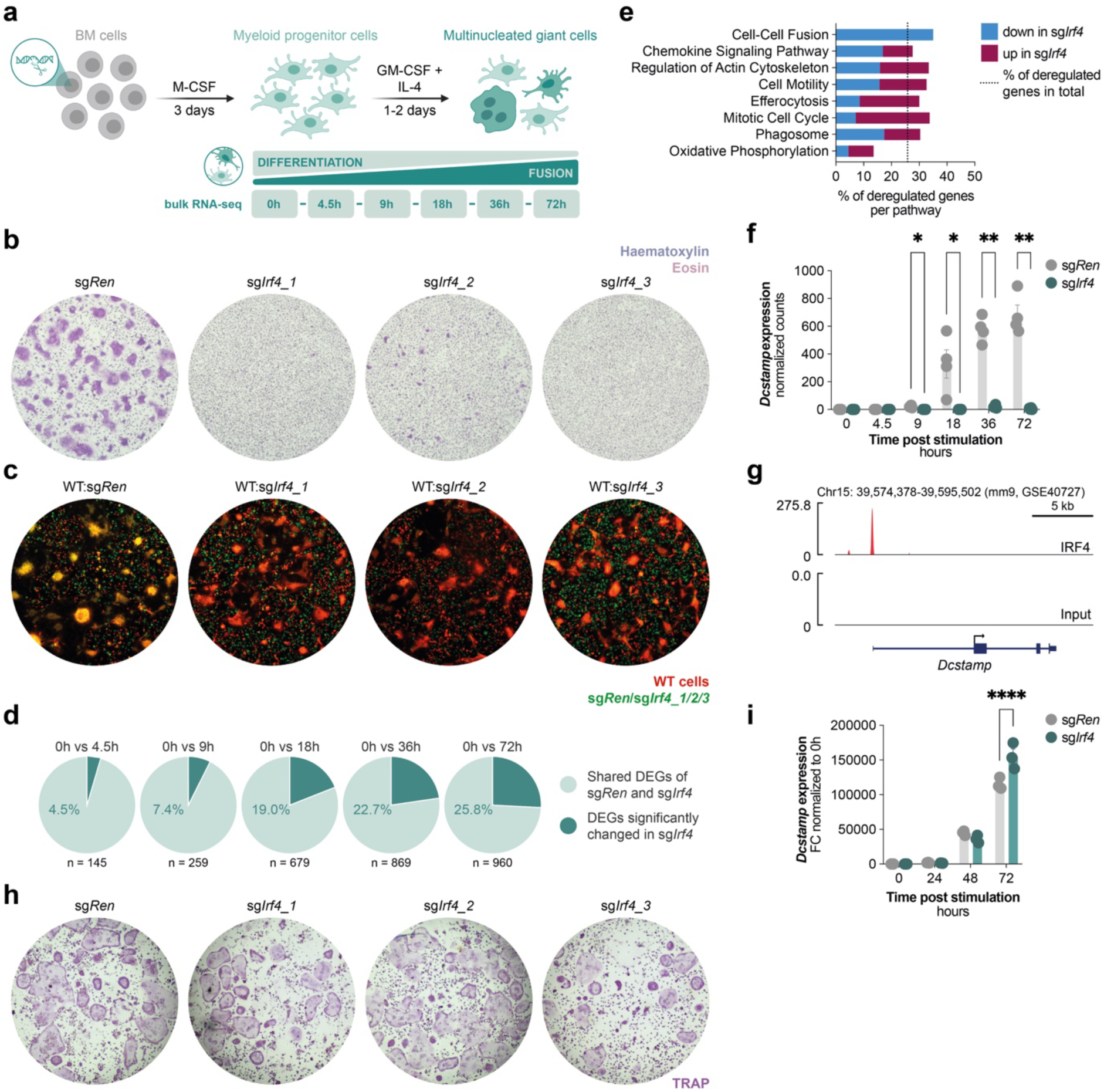
IRF4 specifically regulates cell fusion in MGCs. **(a)** Schematic overview of the *in vitro* formation assay of MGCs derived from CRISPR/Cas9-mediated HoxB8 knockout cell lines including timepoints for bulk RNA-seq sample generation. **(b)** Representative H&E images of CRISPR/Cas9-mediated control (sg*Ren*) or *Irf4* gene (sg*Irf4_1/2/3*) deleted HoxB8 cells stimulated to form MGCs. **(c)** Representative pictures of co-culture experiments of primary BM cells derived from an Ai14^fl^ *LysM-*cre mouse (red; endogenous tdTomato signal) cultured together with CRISPR/Cas9-mediated control (sg*Ren*) or *Irf4* gene (sg*Irf4_1/2/3*) deleted HoxB8 cells (green; AF647-anti-GFP) at a 1:1 ratio and stimulated with IL-4 and GM-CSF. **(d)** Pie charts depicting the proportion of shared DEGs among sg*Ren* and sg*Irf4_1/3* HoxB8 cells of 0h vs various timepoints post fusion induction in light green, as well as the proportion of significantly changed DEGs in sg*Irf4* conditions (sg*Irf4_1*/*3*) vs the control cells (sg*Ren*) in dark green. **(e)** Graph depicting the percentage of *Irf4* dependent deregulated genes (blue: downregulated in sg*Irf4_1/3* and red: upregulated in sg*Irf4_1/3* compared to sg*Ren* of 0h vs 72h as shown in **d** – dark green proportion of pie chart) of selected pathways that have previously been identified to be associated with a fusion competent cell state (**Figure 1e, Suppl. Figure 1c, Figure 2e** and literature based). **(f)** *Dcstamp* expression levels from bulk RNA-seq dataset of sg*Ren* and sg*Irf4_1* HoxB8 cells at different timepoints post fusion induction. **(g)** ChIP-seq tracks for IRF4 (red) at the *Dcstamp* locus derived from a publicly available dataset. **(h)** Representative TRAP stainings of CRISPR/Cas9-mediated control (sg*Ren*) or *Irf4* gene (sg*Irf4_1/2/3*) deleted HoxB8 cells stimulated with RANKL to induce osteoclastogenesis. **(i)** *Dcstamp* mRNA levels from sg*Ren* and sg*Irf4_1* HoxB8 cells at different timepoints during the course of osteoclast formation. Representative images of over 5 independent experiments including at least 2 technical replicates per experiment. **(c)** Representative images of 3 independent experiments. **(d)** and **(e)** Data represent n=4 technical replicates per genotype (sg*Ren*, sg*Irf4_1,* sg*Irf4_3*). **(f)** Data represent n=4 technical replicates per genotype, but only data from sg*Ren* and sg*Irf4_1* HoxB8 cells are shown. **(h)** Representative images of 3 independent experiments including at least 2 technical replicates per experiment. **(i)** Data represent n=3 technical replicates per genotype, but only data from sg*Ren* and sg*Irf4_1* HoxB8 cells is shown. Data are mean ± SEM, *p < 0.05, **p < 0.01 and ****p < 0.0001 two-way ANOVA post-hoc pairwise comparisons with Bonferroni correction.

To analyze the global impact of *Irf4* deficiency on transcriptional events during MGC differentiation, we performed a time course analysis of bulk RNA-seq data of wildtype and *Irf4* knockout cells after stimulation with IL-4 and GM-CSF. We found that *Irf4* deficient cells initially were similar to wildtype cells with only marginal differences in their transcriptome (**Figure 5d** and **Suppl. Figure 5d**). However, consistent with an induction of IRF4 expression during acquisition of fusion competency, the significantly differentially regulated genes between wildtype and knockout cells increased over time and finally comprised about 26% of all genes regulated during MGC formation after 72 hours (**Figure 5d**). Analyzing the distribution of these differentially regulated genes within the pathways we have identified as important during MGC formation, IRF4 specifically controls genes involved in proliferation, cell motility, efferocytosis and phagocytosis (**Figure 5e** and **Suppl. Figure 5e**). In contrast, the geneset describing oxidative phosphorylation was hardly affected by the loss of *Irf4*, implying that the metabolic adaptations required for successful MGC development are independent of this transcription factor. In line, OCR as well as ECAR were not different between wildtype and *Irf4* deficient cells (**Figure 5e** and **Suppl. Figure 5f**). Most importantly, IRF4 pivotally controls the cell fusion pathway, as all genes that are part of this GO-term and significantly regulated during MGC formation were reduced in *Irf4* deficient compared to wildtype cells (**Figure 5e**). Specifically, *Irf4* knockout cells failed to increase the expression levels of well-established cell fusion genes such as *Dcstamp* and *Ocstamp* which are strongly upregulated in wildtype cells upon fusion induction (**Figure 5f, Suppl. Figure 5e and g**). When analyzing a publicly available Chip-seq dataset of GM-CSF and IL-4 derived and LPS stimulated dendritic cells ^32^, we found evidence for binding of IRF4 to the *Dcstamp* promoter (**Figure 5g**), confirming a direct role of this transcription factor for the induction of *Dcstamp* expression during MGC formation.

Osteoclasts share many features with IL-4 induced MGCs, such as the downregulation of macrophage signature genes and the upregulation of DC-STAMP during their differentiation process ^14, 23, 24^. Additionally, the induction of IRF4 expression upon RANKL stimulation has been suggested in a previous study ^33^. Therefore, we tested whether IRF4 would be a general requirement for myeloid multinuclearization. Importantly, both, the capacity of differentiating towards osteoclasts as well as the concomitant increase in the key fusogen *Dcstamp* were not affected in *Irf4* deficient cells (**Figure 5h and i**), suggesting an IRF4 independent pathway in forming multinucleated osteoclasts. Taken together, we identify IRF4 as an essential transcription factor specifically and uniquely regulating the development of MGCs.

## Discussion

The molecular mechanisms governing MGC formation are poorly understood. While the presence of these cells in various pathologies has been known for centuries, virtually no detailed information about the factors driving their generation apart from the cytokines and receptors used to differentiate them, exist ^12, 13^. In this study, we identify the transcription factor IRF4 as a master regulator of MGC development. IRF4 centrally directs the acquisition of a fusion-competent state in myeloid precursor cells as it pivotally controls the expression of DC-STAMP and OC-STAMP, both known to be indispensable for cell to cell fusion ^14, 34^. The role of IRF4 in MGC differentiation closely resembles the one of NFATc1 in osteoclastogenesis. Both transcription factors are not present in the monocytic or macrophage-like precursor cell state but are induced upon cell type specific stimulation towards fusion competency ^21^. In case of MGCs, the specific stimuli comprise GM-CSF and IL-4, whereas osteoclast generation is initiated by RANKL ^19, 35^. While in osteoclasts the upregulation of DC-STAMP requires NFATc1, in MGCs this key fusion factor induction relies on IRF4 ^14^. However, IRF4 in our *in vitro* experiments does not play any role in osteoclast differentiation and vice versa NFATc1 does not influence MGC formation^12^, thus demonstrating the presence of two distinct pathways leading to multinuclearization of myeloid cells, which differ in their regulatory elements, but converge on mutual downstream gene signatures.

Apart from sharing the same key elements of the fusion machinery, we found that IL-4 induced (pre-)MGCs are also metabolically very similar to (pre-)osteoclasts, as both developmental processes are highly energy demanding and therefore require an increase in ATP generation. As in OCs, both OXPHOS and glycolysis are enhanced and functionally required for the differentiation towards a fusion competent cell state, with OXPHOS carrying most of the weight ^36, 37^. However, bioinformatic analysis and metabolic assays revealed that mitochondrial ATP production was only minimally affected by the loss of IRF4, suggesting that the metabolic adaptations of MGC development are not regulated by this transcription factor. Another similarity of osteoclastogenesis and MGC formation is the downregulation of macrophage identity genes like *Mafb* and *Irf8* prior to cell fusion ^22, 23, 24^, which we could confirm in our bulk as well as scRNA-seq dataset. Interestingly, in *Irf4* deficient cells, expression levels of these genes were not decreased upon stimulation with IL-4 and GM-CSF compared to wildtype cells. In addition to the failure of *Irf4* knockout cells to commit to cellular differentiation programs, they seemed to remain in a more proliferative state as a great proportion of genes involved in the mitotic cell cycle pathway were not downregulated in the absence of IRF4.

To elucidate the cellular origins of MGCs *in vivo*, we employed the *S. mansoni* egg-induced MGC formation system in several fate mapping mouse lines. MGCs in lung granulomas of Ai14^fl^ *Ms4a3*-cre mice ^38^ were uniformly and intensely tdTomato positive. This finding indicates that GMP-derived cells serve as the primary precursors for MGCs in our model. A similar pattern of robust tdTomato positivity was evident in Ai14^fl^ *Cd11c*-cre animals, suggesting involvement of CD11c traced myeloid cells in MGC formation. Alveolar macrophages express CD11c and they have previously been implicated in MGC formation in the context of *Aspergillus fumigatus* allergic airway inflammation, indicating that not only infiltrating, but also tissue resident cells might serve as progenitors for MGCs ^30, 39^. However, the analysis of Ai14^fl^ *Cx3cr1*-cre mice revealed a very distinct picture: MGCs were essentially negative for tdTomato labeling in these animals. Since *Cx3cr1*-cre mediated fate mapping is reported to label alveolar macrophages as well as non-classical monocytes ^40^, this suggests that these cell types contribute only minimally to the generation of MGCs in the context of *S. mansoni* egg induced lung granuloma formation. Other CD11c expressing cells that could be involved in MGC generation *in vivo* comprise specific monocyte and dendritic cell subsets. Indeed, previous work could reveal the capability of these cells to serve as osteoclast progenitors ^20^, suggesting a similar role of these cells in *in vivo* MGC formation. Additionally, we could demonstrate that myeloid cells stimulated with IL-4 and GM-CSF concomitantly upregulate the expression of *Itgax* with *Dcstamp* and *Ocstamp in vitro*. Therefore, the tdTomato positivity of MGCs derived from Ai14^fl^ *Cd11c*-cre mice could also be a consequence of CD11c upregulation during MGC formation *in vivo*. Finally, labeling in Ai14^fl^ *LysM*-cre mice was noticeably weaker compared to that seen in Ai14^fl^ *Ms4a3*-or *Cd11c-*cre animals. This observation implies that only a subset of myeloid cells within the MGC population is derived from *LysM* traced precursors. These findings collectively highlight the predominant role of monocyte progenitor cells and/or classical monocytes in the origin of MGCs in the *S. mansoni* egg-induced lung granuloma formation model with limited input from other myeloid lineages, particularly a negligible contribution of *Cx3cr1* traced cells. Importantly, we could demonstrate the presence of IRF4, our proposed master regulator of MGC formation, in MGCs of different cellular origins in lung sections of various pathologies in mice as well as in human derived cells *in vitro* and MGC^high^ samples of head and neck squamous cell carcinomas *in vivo*.

Functionally, the role of MGCs is poorly defined. Previous studies have demonstrated that MGCs were superior in taking up large particles compared to mononucleated myeloid cells ^6^. In line with this data, in our experiments using *S. mansoni* egg induced granuloma formation, we find that MGCs formed around the egg, presumably trying to encapsulate the large parasite derived foreign particle. Additionally, large MGCs were also present in the periphery of the granuloma, with no apparent direct contact to the *S. mansoni* egg. Of note, many of these MGCs seemed to contain polymorphonuclear granulocytes. We therefore propose efferocytosis of granulocytes as an additional function of MGCs in the setting of lung granulomas and provide confirmatory *in vitro* data demonstrating that MGCs indeed engulf apoptotic neutrophils. Interestingly, bioinformatic analysis revealed that IRF4 substantially regulates genes involved in efferocytosis as well as phagocytosis. Among the most significantly downregulated genes in *Irf4* deficient cells, we could identify Axl, a protein importantly involved in the specific uptake of neutrophils ^41^. Altogether, this suggests that in addition to cell fusion, also MGC functional properties might be controlled by IRF4.

Taken together, our data identify IRF4 as an essential regulator of MGC formation.

## Materials and methods

### Animals

Mice were bred and housed in specific pathogen-free facilities at the Medical University of Vienna and kept in a 12h light cycle, at 21–23 °C and 45–65% humidity. All animal procedures were approved by the local ethics committee of the Medical University Vienna and the Austrian Ministry of Sciences (GZ-2025-0.331.342) and were conducted in strict accordance with Austrian law.

Wildtype CD45.2 C57BL/6J animals (C57BL/6J, Jackson Laboratory, #000664) for *in vitro* assays and *in vivo* granuloma formation model were purchased and bred in-house. Wildtype CD45.1 C57BL/6J animals (B6.SJL-Ptprc^a^ Pepc^b^/BoyJ, Jackson Laboratory, #002014) for mixed *in vitro* BM cultures for scRNA-seq data acquisition were used from in-house breedings. Ai14^fl^ mice (B6.Cg-Gt(ROSA)26Sor^tm14(CAG-tdTomato)Hze^/J, Jackson Laboratory, #007914), *Cd11c*-cre mice (B6.Cg^Tg(Itgax-Cre)1-1Reiz^/J, Jackson Laboratory, #008068), *LysM*-cre mice (B6.129P2-*Lyz2^tm1(cre)Ifo^*/J, Jackson Laboratory, #004781), *Ms4a3*-cre mice (C57BL/6J-*Ms4a3^em2(cre)Fgnx^*/J, Jackson Laboratory, #036382), *Cx3cr1*-cre mice (B6J.B6N(Cg)-*Cx3cr1^tm1.1(cre)Jung^*/J, Jackson Laboratory, #025524) and *Arginase1*-eYFP reporter mice (*Arg1*^tm1Lky^, B6.129S4-*Arg1*^tm1.1Lky^/J, Jackson Laboratory, #0015857) have been purchased from Jackson Laboratory. All cre expressing mouse strains were crossed to the Ai14^fl^ strain to generate the respective fate mapping mouse lines. UBI-GFP mice (C57BL/6-Tg(UBC-GFP)30Scha/J, Jackson Laboratory, #004353) were kindly provided by Christoph Binder (Medical University of Vienna, Austria).

The presence of respective transgenes was confirmed by PCR analysis on DNA from ear or toe biopsies of genetically modified animals using the following primer pairs: *Cx3cr1*-cre: CX3CR1_WT: forward, CCTCAGTGTGACGGAGACAG; CX3CR1_mutant: forward, GACATTTGCCTTGCTGGAC; CX3CR1_common: reverse, GCAGGGAAATCTGATGCAAG; *LysM*-cre and *CD11c-*cre: forward, TCGCGATTATCTTCTATATCTTCAG; reverse, GCTCGACCAGTTTAGTTACCC; *Ms4a3*-cre: Ms4a3_WT: reverse, GAAAGGGGAACAAGCGAAGAT; Ms4a3_mutant: reverse, TTGGCGAGAGGGGAAAGAC; Ms4a3_common: forward AGAGAAATCATCAGGGCAGAAAT; Ai14^fl^: tdTomato_WT: forward,<colcnt=2> AAGGGAGCTGCAGTGGAGTA; reverse, CCGAAAATCTGTGGGAAGTC; tdTomato_mutant: forward, CTGTTCCTGTACGGCATGG; reverse, GGCATTAAAGCAGCGTATCC; Arg1-YFP_WT: forward, AGAGCAAGCACCCCGTTTCTTCTC; Arg1-YFP_mutant: forward, TGAGCAAAGACCCCAACGAGAAGC; Arg1-YFP_common: reverse, GCTGTGATGCCCCAGATGGTTTTC.

### *Schistosoma mansoni* egg induced granuloma formation model

Induction of *S. mansoni* egg induced granuloma formation was performed as previously described ^30, 42^. In brief, 5000 *S. mansoni* eggs in sterile 1xPBS (100μl) were injected intraperitoneally (*i.p.*) for sensitization and 14 days later animals were challenged with an intravenous (*i.v.*) injection of 5000 eggs. Healthy control mice were injected with sterile 1xPBS or saline *i.p.* and *i.v.* instead of *S. mansoni* eggs. Lung tissues were harvested on day 7 post *i.v.* injection unless otherwise indicated.

### Immunohistochemistry staining of lung tissue

Animals were euthanized and transcardially perfused with 1xPBS (unless cardiac puncture for blood drawing was performed). Lung tissue was isolated and fixed in 7.5% buffered formalin solution for up to 24h at 4°C. Afterwards, 2μm paraffin-embedded sections were prepared and stained with haematoxylin and eosin (H&E). In brief, slides were stained for 10min with 1:5 diluted Meyer’s hemalum (Merck, #1.09249.0500), rinsed with distilled water, differentiated in 1% HCl-ethanol and rinsed in tap water for 10 min. Afterwards, slides were stained in eosin working solution (300 ml 1% Eosin Sigma-Aldrich, #318906, 600 ml distilled water, 0.1 ml acetic acid 100%) for 15 sec. Slides were then rinsed with distilled water, 96% ethanol, 100% ethanol, n-Butyl-acetate and mounted with Neo-Mount (Merck, #1.09016).

Lung tissue from *S. mansoni* egg induced granuloma formation model and sections from *Aspergillus fumigatus* conidia challenged mice that were kindly provided by David Voehringer (University Hospital Erlangen and Friedrich-Alexander Universität Erlangen-Nürnberg, Germany) were stained with anti-ARG1 (polyclonal, Invitrogen, #PA5-85267, 1:500), anti-IRF4 (clone E8H3S, Cell Signaling Technology, #62834, 1:100) or anti-tdTomato (polyclonal, SICGEN, # AB8181-200, 1:60.000) primary antibodies using different methods for epitope retrieval: heat induced epitope retrieval (HIER) pH6 using citrate buffer and incubating slides for 20min at 96°C for anti-ARG1, HIER pH9 (Epitope retrieval solution pH9, Eubio, #EB-DEPP-9) for 20min at 96°C for anti-IRF4 and protease induced epitope retrieval (PIER) using 0.5 mg/ml Proteinase K (Roche, #03115852001) in 1xPBS for 5min at 37°C for anti-tdTomato staining.

Endogenous peroxidase was blocked after epitope retrieval by incubating sections in 3% peroxide solution for 10min at RT. Unspecific binding of primary antibodies was reduced by blocking sections in 10% serum of the host species of the secondary antibody for 10min at RT. All primary antibody solutions were prepared in 10% serum in 1xPBS (= blocking buffer) and incubated for 1h at RT followed by two washing steps and incubation in the secondary antibody solution prepared in blocking buffer for 30min at RT. Antibody specific signals were developed using the ELITE ABC-HRP Kit (Vector Laboratories, #PK-6100) in combination with DAB Substrate Kit, Peroxidase HRP (Vector Laboratories, #SK-4100) and incubation for 10min at RT. After staining, slides were rinsed in tap water, 96% ethanol, 100% ethanol, n-Butyl-acetate and finally mounted with Neo-Mount (Merck, #1.09016). Image acquisition was performed using a light microscope (Axioskop 2 MOT Zeiss).

### Immunofluorescence staining and image processing of lung tissue

Mice were euthanized and transcardially perfused with 1xPBS (unless cardiac puncture for blood drawing was performed). Lung tissue was isolated and fixed in 7.5% buffered formalin solution for 6h at RT in the dark. Afterwards tissue was saturated with sucrose by transferring to a 30% sucrose/1xPBS solution for 1-3 days and embedded in OCT cryoprotective (Scigen, #4586). 60 µm thick tissue sections were prepared and washed two times in 1xPBS with constant shaking for 10min. Subsequently, blocking solution consisting of 1xPBS supplemented with 5% normal goat serum (Cell Signaling Technology, #5425) and 0.3% Triton X-100 was added and sections were incubated for 3h at RT on a horizontal shaker. After blocking and permeabilization sections were stained with anti-ARG1 primary antibody (polyclonal, Invitrogen, #PA5-85267, 1:100) diluted in blocking solution with constant shaking at 4°C overnight. After washing (4x in 1xPBS for 10min) lung tissue sections were incubated in secondary antibody solution (Alexa Fluor 647 conjugated goat anti-rabbit IgG, Cell Signaling Technology, #4414, 1:500) diluted in blocking solution on a horizontal shaker at 4°C overnight. On the next day, sections were washed (4x in 1xPBS for 10min) and counter-stained with Hoechst 33342 (Invitrogen, #H3570, 1:2500 in 1xPBS) overnight at 4°C. After two washing steps, sections were transferred to slides and mounted in aqueous mounting medium Aquatex (Merck, #10856200500). Images of lung tissue sections were acquired using a spinning disk microscope (Olympus IXplore SpinSR with SoRa Super Resolution Spinning Disk) and processed (adjusting brightness and contrast) and cropped using ImageJ and Adobe Photoshop 2025 (V 26.11).

### Protein isolation and multiplex cytokine analysis of lung tissue

Immediately after harvesting the lung, pieces of different lung lobes where cut, weighed and snap frozen in liquid nitrogen. For protein isolation, frozen lung samples were homogenized in 10μl NP-40 lysis buffer supplemented with protease inhibitors PMSF, PI and SO (all 1:100) per 1mg of tissue using a Precellys tissue homogenizer (Bertin Technologies) (1×15sec at 5800rpm, rest on ice, repeat 1×15sec at 5800rpm). After incubation on ice for 30min, the lung homogenate was centrifuged at 10.000g for 5min at 4°C. The supernatant was transferred into a fresh tube and stored at −80°C until further cytokine detection using a customized LEGENDplex bead-based immunoassay kit (BioLegend).

### Murine MGC and osteoclast culture

Hematopoietic stem cells of the bone marrow (BM cells) were isolated by flushing murine femurs and tibiae with sterile 1xPBS (Gibco, #14190-094). Cell suspension was filtered through a 40µm cell strainer (Falcon, # 352340) and erythrocyte lysis was performed by incubating cells in erythrocyte lysis buffer (0.15M NH_4_Cl, 10mM KHCO_3_, 0.1mM Na_2_EDTA, pH: 7.2 – 7.4) for 5min. Lysis was stopped by adding 1xPBS (10x volume of erythrocyte lysis buffer) and cell suspension was centrifuged at 350g for 5min at RT. Subsequently cells were resuspended in complete MEMα (Gibco, #32561037) containing 5% Pen-Strep (Gibco, #15140122) and 10% fetal bovine serum (Gibco #10082-147). For MGC differentiation assays 450.000 cells per well of a 48-well plate (Thermo Fisher Scientific, #150687) supplemented with 100ng/ml M-CSF (R&D Systems, #416) were incubated for 3 days. After 3 days, medium was changed to complete MEMα supplemented with 50ng/ml IL-4 (R&D Systems, #404) and 50ng/ml GM-CSF (R&D Systems, #415) for another 1-2 days. Inhibitors of the electron transport chain or glycolysis blocking agent 2-Deoxy-D-glucose (2-DG) as well as the respective solvents were added at the same time at the following concentrations: 50nM Oligomycin (MedChemExpress, #HY-N6782, 10mM stock in 100% ethanol), 10nM Antimycin A (Cell Signaling Technology, #33357, 15mM stock in DMSO), 200nM Rotenone (MedChemExpress, #HY-B1756, 10mM stock in DMSO), 100μM 2-Thenoyltrifluoroacetone (TTFA, MedChemExpress, #HY-D0190, 100mM stock in DMSO) and 200μM 2-DG (Sigma-Aldrich, #D8375, 1M stock in 1xPBS). For glucose-free medium, MEMα powder without nucleosides, glucose, pyruvate, vitamin C, amino acids (Genaxxon, #C4002c.005) was dissolved in MiliQ and supplemented with sodiumbicarbonate (Sigma-Aldrich, #S-5761), all amino acids, vitamin C and folic acid used in the formulation of Gibco MEMα as well as with 5% Pen-Strep (Gibco, #15140122) and 10% dialyzed fetal bovine serum (Gibco, #26400044). To rescue the glucose-free condition, 5.5mM glucose was added at the time of fusion induction with IL-4 and GM-CSF.

For live-cell time-lapse microscopy hematopoietic stem cells of UBI-GFP and Ai14^fl^ *Ms4a3*-cre or Ai14^fl^ *LysM*-cre mice were isolated and processed as described above. A mixture of equal amounts of GFP^+^ and tdTomato^+^ cells (in total 400.000 cells) were seeded in a 48-well plate in complete MEMα supplemented with 100ng/ml M-CSF. After 3 days, cells were washed once with complete MEMα before stimulation with either 30ng/ml M-CSF or 50ng/ml IL-4 and 50ng/ml GM-CSF. Pictures of cells were taken every 30-60min for 2 days using an Incucyte S3 Live-Cell Analysis System (Sartorius) with spectral unmixing settings of 60% of red signal contributes to green signal to exclude spillover of tdTomato signal into the green channel. Green + red area per image normalized to values at the start of the assay (0d0h0m) was determined using the Incucyte2024A Software (Sartorius), exported as a Microsoft Excel file and graphed in GraphPad Prism 10 (GraphPad Software). Representative images and movies with adjusted brightness and contrast were generated using the Incucyte2024A Software (Sartorius).

For co-culture experiments of primary murine BM cells with HoxB8 cells, BM cells of Ai14^fl^ *Ms4a3*-cre or Ai14^fl^ *LysM*-cre were isolated and processed as described above. A mixture of equal amounts of GFP^+^ HoxB8 cells (sg*Ren* and sg*Irf4_1/2/3*) and tdTomato^+^ primary murine BM cells (450.000 cells in total) were seeded in a 48-well plate. After 3 days, cells were washed once with complete MEMα before stimulation with IL-4 and GM-CSF for another 1-2 days.

For MGC and osteoclast assays with HoxB8 precursor cells ^43^, HoxB8 cells were washed two times with 1xPBS to remove β-estradiol and GM-CSF and cultured in complete MEMα supplemented with 100ng/ml M-CSF (R&D Systems, #416) at a density of 1×10^6^ cells/ml for MGC (120μl for 96-well plate (Thermo Fisher Scientific, #167008), 300-350μl cell suspension for 48-well plate, 1.1ml for 12-well plate (Thermo Fisher Scientific, #150628) and 2.5×10^5^ cells/ml for osteoclast generation (100μl of cell suspension for 96-well plate, 1ml per 12-well plate). After 3 days medium was changed to complete MEMα supplemented with either 50ng/ml IL-4 (R&D Systems, #404) and 50ng/ml GM-CSF (R&D Systems, #415) or 30ng/ml M-CSF and 50ng/ml RANKL (R&D Systems, #462) for another 2-3 days.

MGC assays were stopped by fixing cells in 3.7% formaldehyde solution in 1xPBS (Sigma-Aldrich, #F8775) for 10min at RT. After washing the cells two times with MiliQ, they were either stained with haematoxylin (Merck, #1.09249.0500) and eosin (Sigma-Aldrich, #318906) or Giemsa staining solution (Sigma-Aldrich, #1092040100). MGCs were defined as cells containing ≥ 3 nuclei. Osteoclast assays were fixed and subsequently stained with TRAP staining solution (Sigma-Aldrich, #387A, Leukocyte Phosphatase Staining Kit). Osteoclasts were defined as TRAP positive multinucleated cells (≥ 3 nuclei). Representative images were acquired using an inverted microscope for cell culture (Olympus CKX53 equipped with an EP50 camera) and further cropped and processed using Adobe Photoshop 2025 (V 26.11).

### Human monocyte-derived MGC cultures

For human MGC assays, CD14^+^ monocytes were sorted from peripheral blood mononuclear cells of healthy donors after density gradient centrifugation and cultured in complete MEMα supplemented with 50ng/mL human M-CSF (R&D systems, #216-MC) or 10μg/ml Concanavalin A (eBioscience, #00-4978-93) at 120.000 cells per 96-well plate or 3.6×10^6^ cells per 1-well chamber slide (Thermo Fisher Scientific, #177372). After 3 days, medium was changed with complete MEMα supplemented with either 50ng/ml human M-CSF and 50ng/ml human IL-4 (R&D systems, #204) or 10μg/ml Concanavalin A (eBioscience, #00-4978-93) or 20ng/ml human GM-CSF (R&D systems, #215) and 10ng/ml human IFNy (PeproTech, #300-02) for another 4 days with one medium change on day3.

### CRISPR/Cas9-mediated gene deletion in HoxB8 cells

HEK293T/17 cells were purchased from ATCC (#CRL-11268) and maintained in DMEM (Gibco, #31966047) completed with 1% Pen-Strep (Gibco, #15140122) and 10% fetal bovine serum (Sigma-Aldrich, #F7524). Immortalized ER-HoxB8 precursors (HoxB8 cells) were provided by Kodi S. Ravichandran (University of Virginia, USA) and were generated as previously described ^44^ from mice with constitutive expression of Cas9 and GFP (Jackson Laboratory, # 026179). HoxB8 cells were maintained in RPMI (Gibco, #61870044) completed with 1% Pen-Strep (Gibco, #15140122), 10% fetal bovine serum (Sigma-Aldrich, #F7524), 1mM Sodium pyruvate (Sigma-Aldrich, #P2256, dissolved in 1xPBS) and MEM non-essential amino acid solution (Thermo Fisher Scientific, #11140050, 1:200) and supplemented with 20ng/ml GM-CSF (R&D Systems, #415) and 1µM β-estradiol (Sigma-Aldrich, #E2758).

CRISPR/Cas9-mediated knockout cell line generation was performed using LentiGuide-Puro (Addgene plasmid #52963). sgRNAs were designed using the Broad Institute sgRNA design tool (https://portals.broadinstitute.org/gppx/crispick/public). The control sgRNA targeting *Renilla luciferase* (sg*Ren*) has been described previously^45^. Cloned oligonucleotides were as follows (5’ to 3’ orientation): *Irf4* sgRNA #1, forward (F): CACCGCAAGCAGGACTACAATCGTG, reverse (R): AAACCACGATTGTAGTCCTGCTTGC; *Irf4* sgRNA #2, forward (F): CACCGGCAGCCTTCAGGGCTCGTCG, reverse (R): AAACCGACGAGCCCTGAAGGCTGCC; *Irf4* sgRNA #3, forward (F): CACCGGGATATCTCTGACCCATACA, reverse (R): AAACTGTATGGGTCAGAGATATCCC; sgRNA oligos with overhangs were annealed using T4 DNA ligase (Thermo Fisher Scientific, #10723941) and ligated into LentiCRISPRv2 construct (Addgene plasmid, 52961) digested with BsmBI-v2 restriction enzyme (New England BioLabs, # R0739S) and amplified in chemically competent (Zymo Research) Stbl3 Escherichia coli cultures (Thermo Fisher Scientific). For lentiviral gene transduction, HEK293T/17 cells were transfected with polyethylenimine (Polysciences Europe, #23966) using the respective lentiviral vector and the packaging plasmids psPAX2 (Addgene plasmid, 12260) and pMD2.G (provided by D. Trono). Medium was changed every day and the cell supernatants from day3 and day4 post transfection were collected, filtered through a 0.45µm polyethersulfone filter (Cytiva, #11344664) and pooled. The supernatants were concentrated by adding a 4x concentrator solution (40% PEG-8000 and 7% NaCl in 1xPBS, pH-adjusted to 7.2 and sterile filtered through a 0.2µm filter) and incubation at 4°C on a rolling shaker (60rpm) for 4.5h, before centrifugation at 3200g for 80min at 4°C. The pellet was dissolved in empty RPMI (= concentrated viral supernatant) and stored at −80°C. Concentrated viral supernatant was diluted in RPMI to a final concentration of 4×10^7^ IU/ml as determined by qPCR using the qPCR Lentivirus Titration Kit (ABM, #LV900) and supplemented with 20ng/ml GM-CSF, 1µM β-estradiol, 8µM cyclosporine H (AdooQ Biosciences, #A15435) and 1mg/ml LentiBoost P (Sirion Biotech, #SB-A-LF-901) before addition to stably Cas9 expressing HoxB8 cells. Spinfection was performed at 800g for 45min at RT. At day1 post infection, medium was changed to complete HoxB8 RPMI and at day2 successfully infected HoxB8 cells were selected by adding 8µg/ml Puromycin (Sigma-Aldrich, #P8833). Cells were maintained on selection for 2 days. The knockout efficiency was determined using western blotting.

### Efferocytosis assay

Neutrophils were isolated from BM cells after performing erythrocyte lysis by positive selection with APC-conjugated anti-Ly6G (clone REA526|1A8, Miltenyi Biotec, #130-120-803) and anti-APC MicroBeads (Miltenyi Biotec, #130-090-855) following the manufacturer’s instructions. Afterwards apoptosis of neutrophils was induced by culturing them in RPMI (Gibco, #61870044) completed with 1% Pen-Strep (Gibco, #15140122) and reduced serum (2-5% fetal bovine serum (Sigma-Aldrich, #F7524)) for 24h ^41^. Apoptotic levels were verified using FITC Annexin V (BioLegend, # 640906, 1:20) staining in Annexin V Binding Buffer (BioLegend, # 422201) and subsequent DAPI staining (Sigma-Aldrich, #D9542, 1mg/ml stock used 1:1000). To be able to track apoptotic neutrophil uptake, 1×10^6^ neutrophils/ml were labelled with pHrodo Red Cell Labelling Kit for Incucyte (Sartorius, #4649, 1:10.000 in 1xPBS) for 20min at RT in the dark. After one washing step with 1xPBS, labelled apoptotic neutrophils were incubated with either previously generated primary mouse MGCs (1 day post fusion induction) or HoxB8 derived MGCs (2-3 days post fusion induction) in a 3:1 or 2:1 ratio (apoptotic cells:MGCs). As positive and negative technical staining controls, apoptotic neutrophils were cultured alone in citrate buffer pH4.5 or complete MEMα, respectively. To detect potential autofluorescence signals of MGCs, wells without labelled apoptotic neutrophils were tracked as well. Pictures of cells were taken every 30-60min for up to 24h using an Incucyte S3 Live-Cell Analysis System (Sartorius) with phase and red fluorescence channel detection. Representative images and movies with adjusted brightness and contrast were generated using the Incucyte2024A Software (Sartorius) and Adobe Photoshop 2025 (V 26.11).

### Immunohistochemistry and immunofluorescence staining of *in vitro* assays

For immunohistochemistry staining, fixed MGC cultures were washed once with ice-cold 1xPBS before permeabilization and peroxide blocking in 0.3% H_2_O_2_ solution in methanol (stored at −20°C) by incubation for 30min on ice on a horizontal shaker. After two washing steps with 1xPBS (5min each), cells were incubated in blocking buffer consisting of 5% normal goat serum (Cell Signaling Technology, #5425) in 1xPBS for 1-2h at RT. Afterwards cells were stained with anti-IRF4 (clone E8H3S, Cell Signaling Technology, #62834, 1:200 in blocking buffer for murine cells or 1:100 in blocking buffer for human cells) at 4°C overnight with constant shaking. On the next day, cells were washed three times with 1xPBS, followed by incubation with the secondary goat anti-rabbit horseradish peroxidase-linked secondary antibody of the Elite ABC-HRP Kit (Vector Laboratories, #VECPK-6101) in blocking buffer for 1h at RT. After two washing steps in 1xPBS, the antibody specific signal was developed by first incubating cells in ABC reagent and after another two washing steps in 1xPBS, subsequent incubation in the peroxidase substrate solution (DAB Substrate Kit, Vector Laboratories, # VECSK-4100) until the desired staining intensity was visible. Cells processed in the same way except incubation with primary antibody served as negative staining controls. In some experiments counterstaining was performed using 1:5 diluted Meyer’s hemalum (Merck #1.09249.0500) for 5min, rinsed first with distilled water and then with tap water. Representative images were taken using an inverted microscope for cell culture (Olympus CKX53 equipped with an EP50 camera) and further cropped and processed using Adobe Photoshop 2025 (V 26.11).

For immunofluorescence staining of fixed MGC (co-)cultures cells were washed two times in 1xPBS before permeabilization and blocking in blocking solution consisting of 5% normal goat serum (Cell Signaling Technology, #5425) and 0.3% TritonX-100 in 1xPBS for 1h at RT. Afterwards endogenous GFP signal of HoxB8 cells was enhanced by staining with anti-GFP antibody (Abcam, #ab13970, 1:400 for overnight incubation or 1:250 for incubation at RT for 1h) diluted in blocking buffer. On the next day, cells were washed three times with 1xPBS, followed by incubation in the secondary antibody solution (Alexa Fluor 647 conjugated goat anti-chicken IgG, Invitrogen, #A32933TR, 1:400) diluted in blocking buffer for 1h at RT. For additional IRF4 staining, cells were washed three times in 1xPBS, again blocked and permeabilized in blocking solution and first incubated with anti-IRF4 antibody solution (clone E8H3S, Cell Signaling Technology, #62834, 1:200 in blocking buffer) at 4°C overnight, followed by washing steps and incubation with the secondary antibody solution (Alexa Fluor 555 conjugated goat anti-rabbit IgG, Thermo Fisher Scientific, #A21428, 1:500) diluted in blocking buffer for 1h at RT. For some experiments counterstaining was performed using Hoechst 33342 (Invitrogen, #H3570, 1:500 in 1xPBS) for 10min. Representative images were acquired using a live imaging microscope (Olympus IX83 or Olympus IX71 equipped with an ANDOR iXon Life 888 camera) and further cropped and processed using ImageJ and Adobe Photoshop 2025 (V 26.11).

For immunofluorescence staining of fixed MGCs generated from *Arginase1*-eYFP reporter mice, cells were washed two times in 1xPBS before permeabilization in ice-cold methanol (stored at −20°C) for 10min on ice. Subsequently, cells were blocked in blocking solution consisting of 5% normal goat serum (Cell Signaling Technology, #5425) and 0.3% TritonX-100 in 1xPBS for 1h at RT. Afterwards staining procedure for enhancing the endogenous YFP signal was continued as outlined above except using an Alexa Fluor 488 conjugated goat anti-chicken IgG (Invitrogen, #A11039, 1:400) secondary antibody diluted in blocking buffer for 1h at RT. Representative images were acquired using an Olympus IX71 microscope equipped with an ANDOR iXon Life 888 camera and further cropped and processed using ImageJ and Adobe Photoshop 2025 (V 26.11).

### RNA isolation and quantitative PCR of tissue und cells

Total RNA of lung pieces of different lung lobes (snap frozen in liquid nitrogen directly after harvest) was extracted by tissue homogenization in TRIzol reagent (Invitrogen, #12034977) using a Precellys tissue homogenizer (Bertin Technologies) and RNA clean-up post isolation with chloroform using the Monarch RNA Clean-up kit (NEB, #T2040). Reverse-transcription was performed using High-Capacity cDNA Reverse Transcription Kit (Applied Biosystems, #4368814). Luna Universal qPCR Master Mix (NEB, #M3003E) was used for the quantitative PCR reaction. Postamplification melting curve analysis and water controls were included to ensure absence of primer dimers. To obtain sample-specific ΔCt values, normalisation to hypoxanthine phosphoribosyltransferase 1 (*Hprt*) (for data with murine BM cells and HoxB8 cells) and beta-actin (*Actb*) (for data with lung tissue) within each sample was performed. For data with human sorted CD14^+^ monocytes normalisation to *HPRT* within each sample was performed. Data are shown as fold change, where 2−ΔΔCt values were calculated (ΔΔCt = ΔCt treatment − ΔCt control). Real-time PCR was performed using the following primers for murine genes: *Hprt*: 5′-*CGCAGTCCCAGCGTCGTG*-3′ and 5′-*CCATCTCCTTCATGACATC* TCGAG-3′; *Actinb*: 5’-*TGTCCACCTTCCAGCAGATGT*-3’ and 5’-*AGCTCAGTAACAGTCCGCCTAGA*-3’; *Dcstamp*: 5′-*TCCTCCATGAACAAACAGTTCCAA*-3′ and 5′-*AGACGTGGTTTAGGAATGCAGCTC*-3′; *Irf4*: 5′-*TGCAAGCTCTTTGACACACA*-3′ and 5′-*TGGAAACTCCTCACCAAAGC*-3′. Real-time PCR was performed using the following primers for human genes: *HPRT*: 5’-*CTCATGGACTGATTATGGACAGGAC*-3’ and 5’-*GCAGGTCAGCAAAGAACTTATAGCC*-3’; *IRF4*: 5′-*AGCAGTTCTTGTCAGAGC*-3′ and 5′-*GTTCTACGTGAGCTGTGATG*-3′; *DCSTAMP*: 5′-*CTTGCCAGGGTTTGAGGTTCAC*-3′ and 5′-*GGGATACAGTTGGGTTCAAACAC*-3′.

### Western blot

Cells were washed with ice-cold PBS before addition of RIPA lysis buffer supplemented with protease inhibitors included in the kit (Santa Cruz, #24948). After incubation for 5min on ice with constant shaking, cells were scraped and transferred to a 1.5ml reaction tube. Cell lysis and protein extraction was continued by incubation on ice for 15min with three vortexing steps in between. The homogenate was finally cleared by centrifugation at 4°C for 15min at 16.000g and the supernatant containing the protein fraction was recovered. Protein concentration was determined using the Pierce BCA Protein Assay Kit (Thermo Fisher Scientific, #23225). A total of 15µg of protein was resolved by SDS-PAGE and transferred to PVDF membranes (GE Healthcare, #10600023). Membranes were blocked in TBS-Tween (0.1%) containing 5% skim milk powder (Sigma-Aldrich, #70166) and incubated with constant agitation with primary antibody solution at 4°C overnight. The following antibodies were used: anti-IRF4 (clone E8H3S, Cell Signaling Technology, #62834, 1:1000 in blocking buffer) and anti-GAPDH (clone D16H11, Cell Signaling Technology, #5174, 1:2000 in blocking buffer). The incubated membranes were washed three times for 5min in TBS-Tween (0.1%) and probed with a goat anti-rabbit horseradish peroxidase-linked secondary antibody (Cell Signaling Technology, #7074, 1:5000 in blocking buffer). Antigen-specific binding of antibodies was detected with WesternBright Sirius HRP substrate (Advansta, #541020). Western blots were assessed by an area density analysis using the Vision Works Software (Analytik Jena).

### Real-Time cell metabolic analysis

Oxygen consumption rate (OCR) and extracellular acidification rate (ECAR) of primary mouse BM cells were measured on a Seahorse XF HS mini Analyzer (Agilent) using the Seahorse XFp Real-Time ATP Rate Assay kit (Agilent, #103591-100). In brief, 30.000 BM cells were seeded per well in a XF HS mini plate using complete MEMα supplemented with 100ng/ml M-CSF (R&D Systems, #416). After 3 days, cell medium was changed to complete MEMα supplemented with 50ng/ml IL-4 (R&D Systems, #404) and 50ng/ml GM-CSF (R&D Systems, #415). At different timepoints post fusion induction (0h, 6h, 12h and 24h) medium was changed to XF RPMI medium (Agilent, #103576-100) supplemented with Sodium pyruvate, Glutamine and Glucose (concentrations equal to MEMα) and plates were incubated in a non-CO_2_ incubator for 1h for equilibration before measurements. During the measurements, 1.5µM oligomycin and 500nM rotenone/antimycin A were subsequently injected. OCR and ECAR of HoxB8 cells were measured on the same Seahorse Analyzer using the Seahorse XFp Cell Mito Stress test kit (Agilent # 103010-100) according to the manufacturer’s instructions. In brief, 15.000 HoxB8 cells were seeded per well in a XF HS mini plate using complete MEMα supplemented with 100ng/ml M-CSF. After 3 days, cell fusion was induced by medium change to complete MEMα containing 50ng/ml IL-4 and 50ng/ml GM-CSF. After 24h, medium was changed to XF DMEM medium (Agilent, #103575-100) supplemented with 1mM Sodium Pyurvate, 2mM Glutamine and 10mM Glucose and plates were kept in a non-CO_2_ incubator for 1h for equilibration before measurement. During the measurements, 1.5µM oligomycin, 1.5µM FCCP and 500nM rotenone/antimycin A were subsequently injected. Raw data were analyzed using Seahorse Analytics Online Software system, exported as a Microsoft Excel file and graphed in GraphPad Prism 10 (GraphPad Software).

### Bulk transcriptomics and analysis

For bulk RNA-seq data of primary murine BM cells total RNA was extracted from 1.5×10^6^ seeded cells (12-well plate) using TRIzol reagent (Invitrogen, #12034977) and the Monarch RNA Clean-up kit (NEB, #T2030). For bulk RNA-seq data of HoxB8 cells total RNA was extracted from 1.1×10^6^ seeded cells (12-well plate) using the above-described reagents and methods.

NGS libraries were prepared using the QuantSeq 3’ mRNA-Seq FWD library preparation protocol (Lexogen GmbH, Vienna, Austria) and sequenced on a HiSeq 3000 instrument (Illumina, San Diego, CA, USA) following a 50-base-pair, single-end recipe. Raw data acquisition and base calling were performed with HiSeq Control Software (HCS HD 3.4.0.38) and Real-Time Analysis (RTA 2.7.7), respectively. FASTQ files were adapter- and quality-trimmed using Trim Galore! (v0.4.4), then aligned to the mouse reference genome (GRCm38) with STAR (v2.5.2; ^46^). Differential expression was analyzed in R (v4.0.4) using DESeq2 (v1.30.1) and genes with an absolute shrunken fold change > 1.5 and FDR < 0.05 were considered significant ^47^. Soft clustering of longitudinal gene expression profiles was performed on VST-transformed expression values which were imported into R using the Mfuzz package (table2eset function), gene-wise standardized (standardise function), and soft-clustered (mfuzz function; c=4, m=1.7). Data were visualized in MA plots (R package ggpubr), volcano plots (VolcaNoseR^48^), heatmaps (ClustVis ^49^) and pie charts. For pie charts DEG lists (FDR < 0.05) of sg*Ren* conditions (0h vs xh) were generated and overlayed with the expression data from sg*Irf4_1* and sg*Irf4_3* conditions (0h vs xh). Differences in the shrunkLog2 FC values between knockout and wildtype data were calculated and only genes that passed the log2FC difference cut-off of < −1 or > 1 were considered to be differentially regulated in the *Irf4* knockout condition versus wildtype cells (dark green part of piecharts) at the respective timepoint comparisons. For transcription factor analysis data was overlayed with the transcription factor data base TFDB v4.0 ^50^. Functional enrichment analysis was performed using the GO Biological Process database implemented in g:Profiler (version: e111_eg58_p18_b51d8f08,^51^ with g:SCS multiple testing correction method applying significance threshold of 0.05 and using all genes for which counts were detectable in the bulk RNA-seq dataset as background gene list.

### Single cell transcriptomics and analysis

Hematopoietic stem cells of CD45.1 and CD45.2 animals were isolated and processed as described for *in vitro* MGC cultures. A mixture of equal amounts of CD45.1 and CD45.2 cells (in total 1.5×10^6^ cells in total) were seeded per well of a 12-well plate in complete MEMα supplemented with 100ng/ml M-CSF. After 3 days, cells were washed once with 1xPBS before fusion induction with 50ng/ml IL-4 and 50ng/ml GM-CSF at different timepoints (reverse time course experiment). Cells of all conditions (0h, 6h, 12h of stimulation with IL-4 and GM-CSF) were prepared for subsequent 10xGenomics at the same time as follows: First, cells were washed in 1xPBS before incubation in TruStain FcX blocking solution (BioLegend, #101320, 1:100 in 1xPBS) for 10min at 4°C on shaker. CITE-seq antibody solution consisting of TotalSeq-anti-CD45.1 (clone A20, BioLegend, # 110753, final dilution after adding to cells 1:100) and TotalSeq-anti-CD45.2 (clone 104, BioLegend, #109853, final dilution after adding to cells 1:100) in 1% FCS in 1xPBS was added to the cells followed by incubation for 30min at 4°C on a horizontal shaker. After three washing steps in 1% FCS in 1xPBS (5min each at 4°C) cells were detached using pre-warmed (37°C) enzyme free dissociation solution (Sigma-Aldrich, #S-014-B) by incubating cells for 10min at 37°C. Cells were harvested by gently pipetting up and down and pelleted at 250g for 5min at 4°C. Subsequently, cell pellets were resuspended in 200µl of 0.04% BSA in 1xPBS and filtered through a 40µm Flowmi cell strainer (Sigma-Aldrich, #BAH136800040) to generate a single cell suspension without cell clumps. Cell counts and viability were determined using an automated cell counter with trypan blue. Appropriate volumes of 0.04% BSA in 1xPBS were added to each cell suspension to get a cell density of 1000 cells/µl.

Single cell RNA-seq libraries were generated using the Chromium iX Controller and the Next GEM Single Cell 3’ Reagent Kit (v3.1, 10x Genomics) according to the manufacturer’s instructions. The samples were processed individually according to the manufacturer’s protocol, with the antibody-linked barcodes enabling protein detection via sequencing. Libraries were sequenced by the Biomedical Sequencing Facility at the CeMM Research Center for Molecular Medicine of the Austrian Academy of Sciences, using the Illumina NovaSeq 6000 platform. Raw sequencing data was pre-processed and demultiplexed using bcl2fastq (v 2.20.0.422) and Cell Ranger (v 5.0.1, 10x Genomics).

Antibody-linked barcodes were used to potentially enable the identification of early fused cells (CD45.1^+^ CD45.2^+^ cells). However, since no robust double-positive signal could be detected, the antibody-linked barcodes were not used for further analysis. Gene-barcoded matrices from three time points (0h, 6h, 12h) were imported with Seurat (version 4.0.6; Read10X). For each time point we created a Seurat object from the RNA assay (CreateSeuratObject, min.features=200, min.cells=3). Mitochondrial transcript proportion was computed and cells were retained if nFeature_RNA > 200, nFeature_RNA < 2,500, and percent.mt < 10%. After QC, data were normalized (NormalizeData), highly variable genes were selected (FindVariableFeatures), counts were scaled (ScaleData), and dimensionality reduction performed by PCA (RunPCA). A shared nearest neighbor graph was constructed on PCs 1–30 (FindNeighbors), clusters were identified at resolution = 0.5 (FindClusters), and visualized with UMAP (RunUMAP).

For multi-sample integration, per-time-point objects were merged and a batch key was recorded in metadata. We used Seurat’s anchor-based workflow: integration anchors were computed on PCs 1–30 (FindIntegrationAnchors), followed by data integration (IntegrateData). The integrated assay was scaled and reduced, neighbors and clusters recomputed on PCs 1–30, clusters were identified at resolution = 0.25, and visualized with UMAP. Differential expression was calculated with pairwise comparisons between cluster–time combinations; statistics used Seurat’s default Wilcoxon rank-sum test (FindMarkers). Functional enrichment analysis was performed using the KEGG database implemented in g:Profiler (version: e113_eg59_p19_f6a03c19^51^) with g:SCS multiple testing correction method applying significance threshold of 0.05 and using a custom background gene list (all genes included in the Seurat object).

### Analysis of publicly available datasets

Publicly available processed 10x Genomics Visium spatial transcriptomics data for head and neck squamous cell carcinoma (HNSCC) were downloaded from ArrayExpress (E-MTAB-14409) in .cloupe format and visualized with Loupe Browser v8 (10x Genomics). Preprocessed murine Irf4 ChIP-seq signal tracks were downloaded from the Gene Expression Omnibus (GEO; GSE40727) as wiggle (.wig) files. Wig files were converted to bigWig using the UCSC utility wigToBigWig with the matching mouse chromosome sizes file, and exported to bedGraph via bigWigToBedGraph. Tracks were visualized with SparK (https://github.com/harbourlab/SparK).

## Data and code availability

The data that support the findings of this study are available from the corresponding authors upon reasonable request. Bulk RNA-seq datasets generated in this study have been deposited in ArrayExpress under accession number E-MTAB-15685. The results published here are in part based on analysis of publicly available spatial transcriptomics and ChIP-seq data deposited under ArrayExpress data set E-MTAB-14409 and GEO data set GSE40727.

## Statistics

Statistical analysis was performed using a two-tailed Student’s t-test for two groups, an ordinary one-way ANOVA followed by Tukey’s multiple comparisons test or a two-way ANOVA followed by Bonferroni’s multiple comparisons test using Prism 10 software (GraphPad, La Jolla, CA). Statistical significance is indicated by *p < 0.05, **p < 0.01, ***p < 0.001, ****p < 0.001. All error bars indicate ±SEM.

## Author contributions

M.H., S.B. and G.S. conceived and designed the study. G.H. developed the computational framework and headed bioinformatic analysis supported by M.H. . M.H. performed and analyzed data for most of the experiments with support from M.Ki., L.M., L.K., M.Ke., P.E., B.N., L.Q.G. and A.S. . L.X.H., T.G., T.W., M.Ki., and L.M. provided important technical input. B.E., A.D., and D.V. provided key resources. M.H. and S.B. wrote the manuscript with important input from O.S. and G.S. . All authors read, revised and approved the final manuscript.

## Acknowledgements

We thank Hans Häcker and Kodi Ravichandran for providing HoxB8 cells. This research was funded by the Christian Doppler Laboratory for Arginine Metabolism in Rheumatoid Arthritis and Multiple Sclerosis as well as the Spezialforschungsbereich (SFB) F83 Immunometabolism to G.S., the FWF (Grant-DOI: 10.55776/PAT5446023) to S.B. and the Deutsche Forschungsgemeinschaft (DFG) grants VO944/11-1 and CRC1181 project A2 to D.V. . M.H. was rewarded a DOC fellowship by the Austrian Academy of Sciences. We thank the Biomedical Sequencing Facility (BSF) at the Research Centre for Molecular Medicine of the Austrian Academy of Sciences (CeMM) for their support in generating scRNA- and bulk RNA-seq data as well as the staff at the Imaging Core facility, Medical University of Vienna, for their assistance in the acquisition of lung tissue images. Schemes depicted in Figures 1, 2, 3 and 5 were created with BioRender.com.

---

**Suppl. Figure 1:**
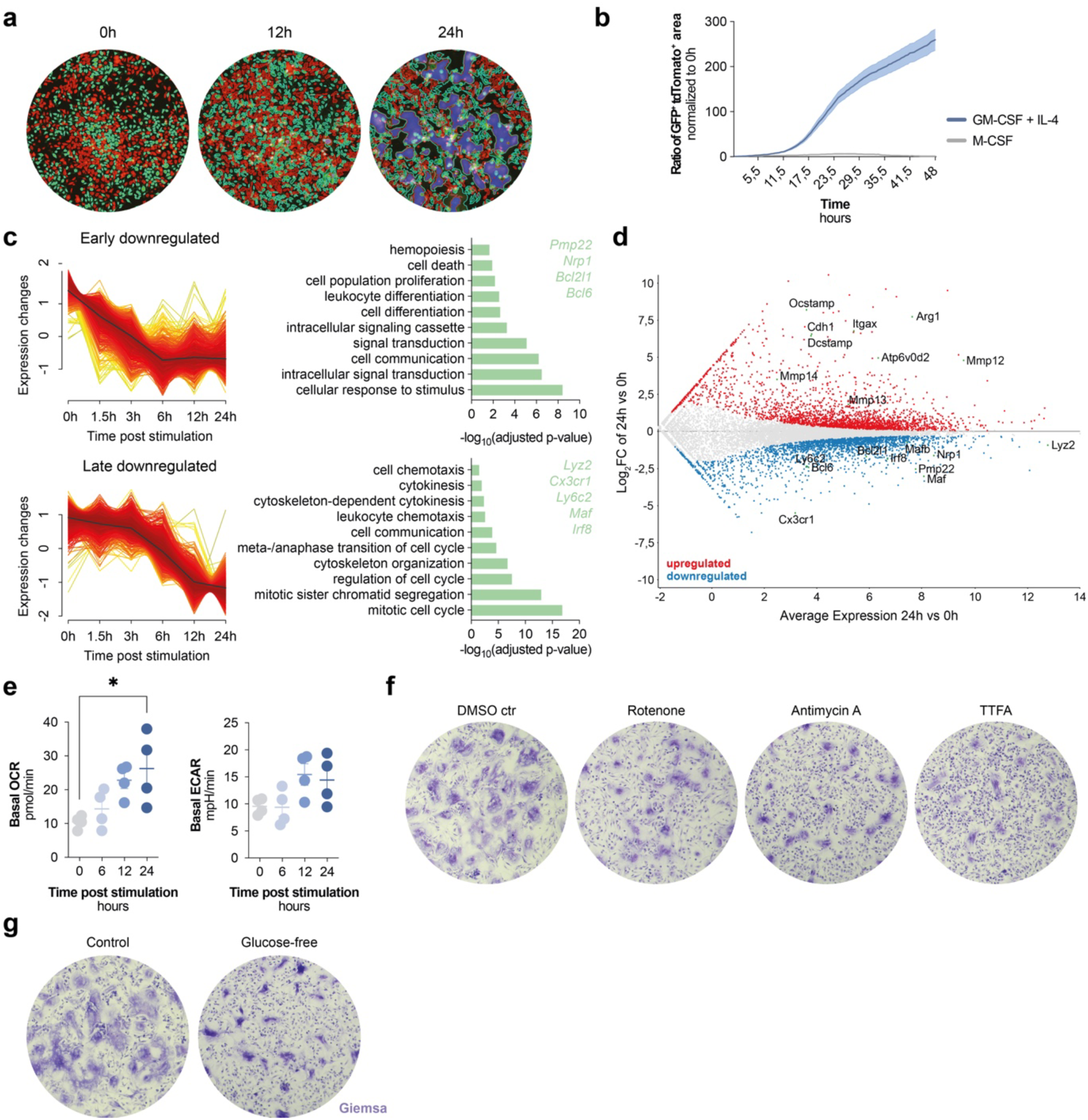
**(a)** Representative images of GFP^+^ (circled in green), tdTomato^+^ (circled in red) and GFP^+^ tdTomato^+^ (labelled in blue) cell area classification using the Incucyte2024A Software (Sartorius) of cells depicted in Figure 1b. **(b)** Representative quantification of the ratio of GFP^+^ tdTomato^+^ area normalized to the baseline values (0h timepoint) of myeloid precursors stimulated either with M-CSF or IL-4 and GM-CSF for up to 48h. **(c)** Unbiased gene expression pattern analysis plots of the 3197 putative master fusion genes shown in Figure 1d and a selection of significantly enriched pathways described by genes included in the respective clusters as determined by ORA based on the GO biological pathway (GO:BP) database. Upper and lower panel represent the early and late downregulated gene trajectories and their respective selections of significantly enriched GO:BP pathways as well as a selection of genes included in these trajectories, respectively. **(d)** MA-plot depicting DEGs between the 0h and 24h timepoint. Up-(red) and downregulated (blue) genes based on adjusted p-value of 0.05. **(e)** Basal OCR and ECAR levels of myeloid cells at various timepoints post fusion induction. **(f)** Representative images of the inhibiting effect of 200nM Rotenone, 10nM Antimycin A and 100µM TTFA on MGC formation of murine BM cells including DMSO solvent control. **(g)** Representative pictures of myeloid precursor cells that were differentiated towards MGCs in the presence or absence of glucose. **(a)** Representative images of in total n=2 biological replicates of 2 independent experiments. **(b)** Data represent n=26-27 images of 3 independent wells (representative data of 2 independent experiments). **(c)** and **(d)** Data represent n=4 biological replicates per timepoint. **(e)** Data represent n=4 biological replicates per timepoint of 2 independent experiments. **(f)** and **(g)** Representative images of n=2-4 biological replicates of up to 2 independent experiments. Data are mean ± SEM, *p < 0.05, one-way ANOVA post-hoc pairwise comparisons with Tukey correction.

**Suppl. Figure 2:**
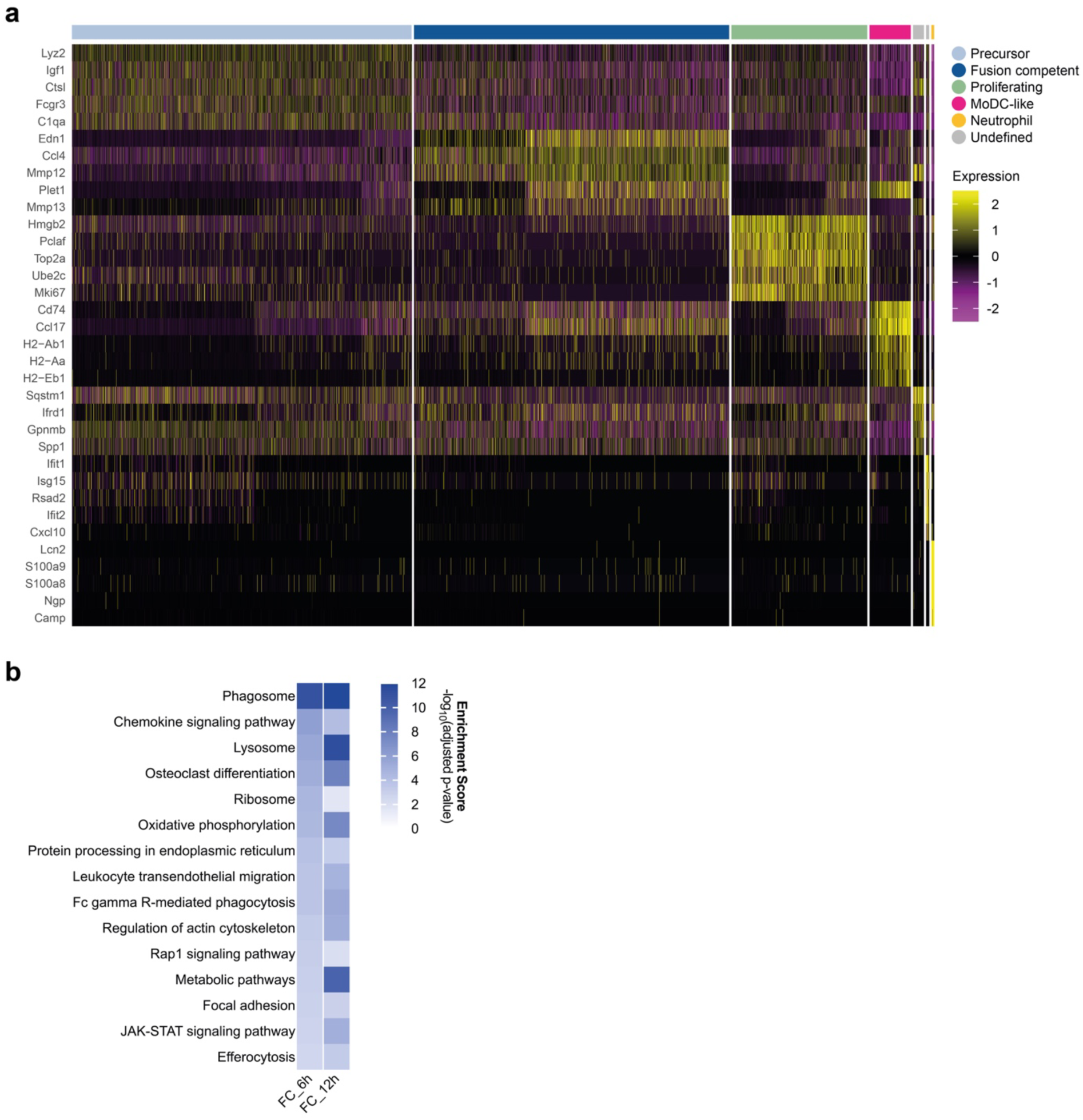
**(a)** Heatmap of top 5 marker genes that were differentially expressed in each cluster. **(b)** Heatmap depicting the enrichment scores of selected significantly enriched pathways of genes differentially expressed between fusion competent cells of either the 6h or 12h timepoint, both compared to precursor cells at the baseline (0h) as determined by ORA based on the KEGG database.

**Suppl. Figure 3:**
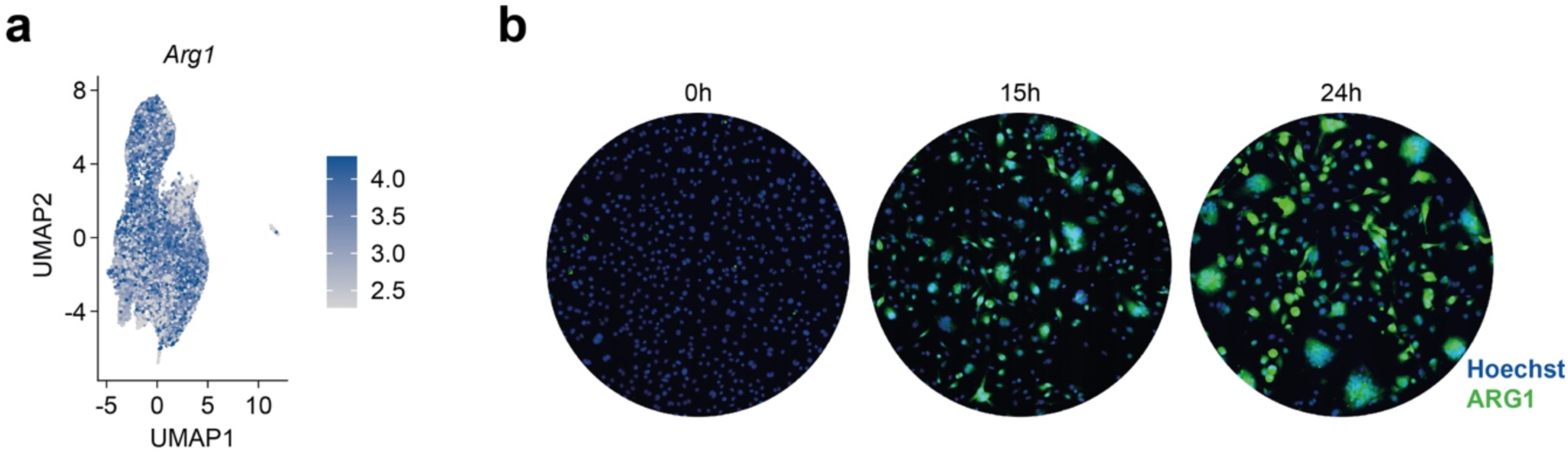
**(a)** The expression pattern of *Arg1* in the UMAP visualization of our scRNA-seq dataset. **(b)** Representative images of myeloid precursor cells derived from an *Arginase1*-eYFP reporter mouse stimulated with IL-4 and GM-CSF for the indicated timepoints and stained with anti-GFP (AF488, green) to enhance the endogenous YFP signal and Hoechst (blue). **(a)** Data represent n=2 pooled biological replicates. **(b)** Representative images of n=2 biological replicates.

**Suppl. Figure 4:**
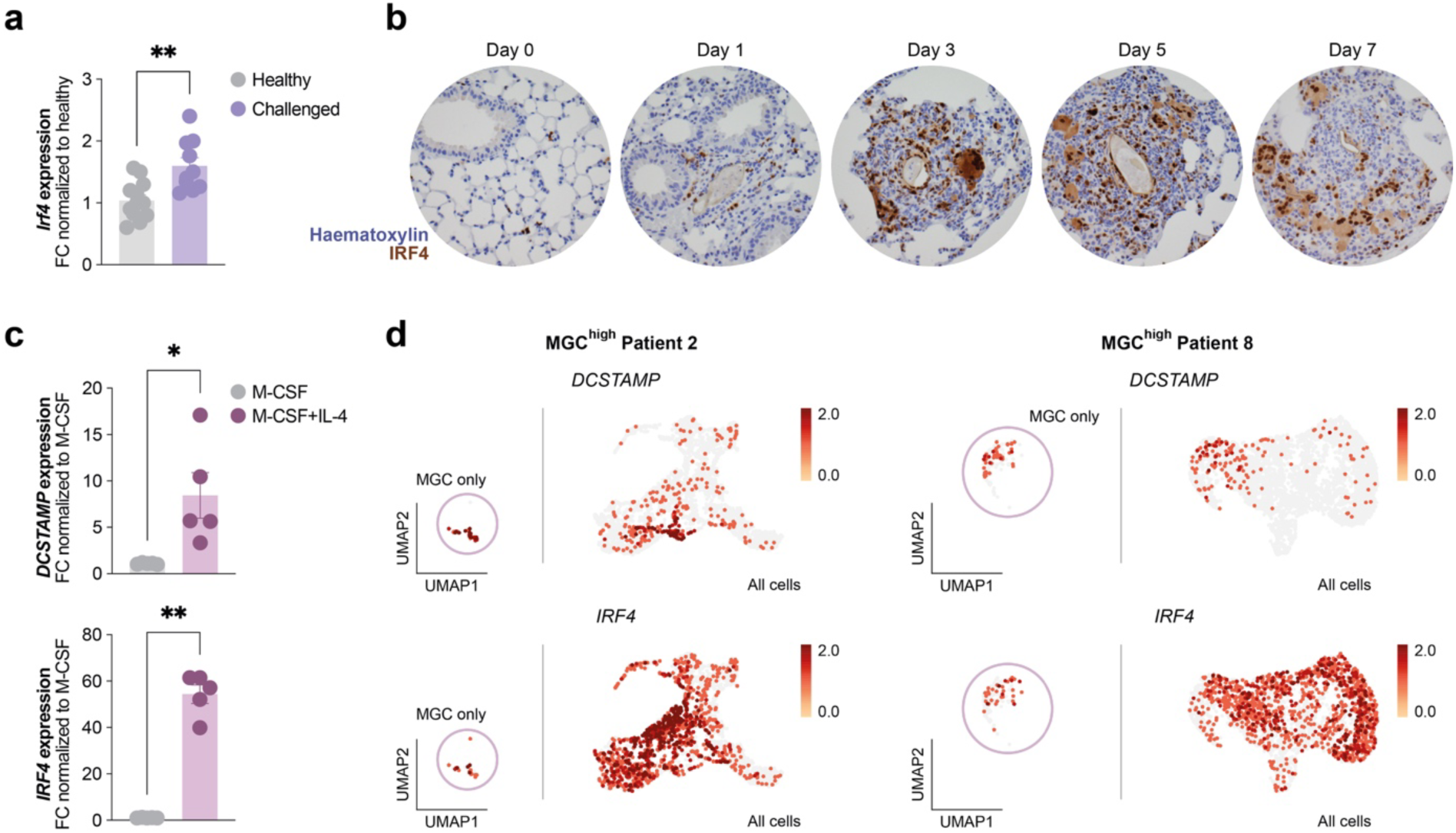
**(a)** Relative mRNA expression of *Irf4* in lung tissue of healthy and challenged animals on day 7 post *i.v.* challenge with *S. mansoni* eggs. **(b)** Representative pictures of IRF4 staining in lungs on various timepoints post *i.v.* challenge with *S. mansoni* eggs. **(c)** *DCTSAMP* and *IRF4* mRNA expression levels of human derived monocytes cultured in the presence of M-CSF alone or in the combination of M-CSF and IL-4 until the end of MGC formation. **(d)** Expression pattern of *DCSTAMP* and *IRF4* of the MGC^high^ Patient 2 and 8 samples derived from a publicly available dataset either depicted for the MGC cluster alone (left) or for all cells together (right) in the UMAP visualization. **(a)** Data represent n=12 healthy and n=10 challenged mice of 3 independent experiments. **(b)** Representative images of n=5 animals for day0, n=3 for day1, n=7 for day3, n=7 for day5, n=6 for day7 post *i.v.* challenge with *S. mansoni* eggs of 2 independent experiments. **(c)** Data represent n=5 individual donors per condition pooled from 5 independent experiments. Data are mean ± SEM, *p < 0.05 and **p < 0.01 unpaired *t*-test.

**Suppl. Figure 5:**
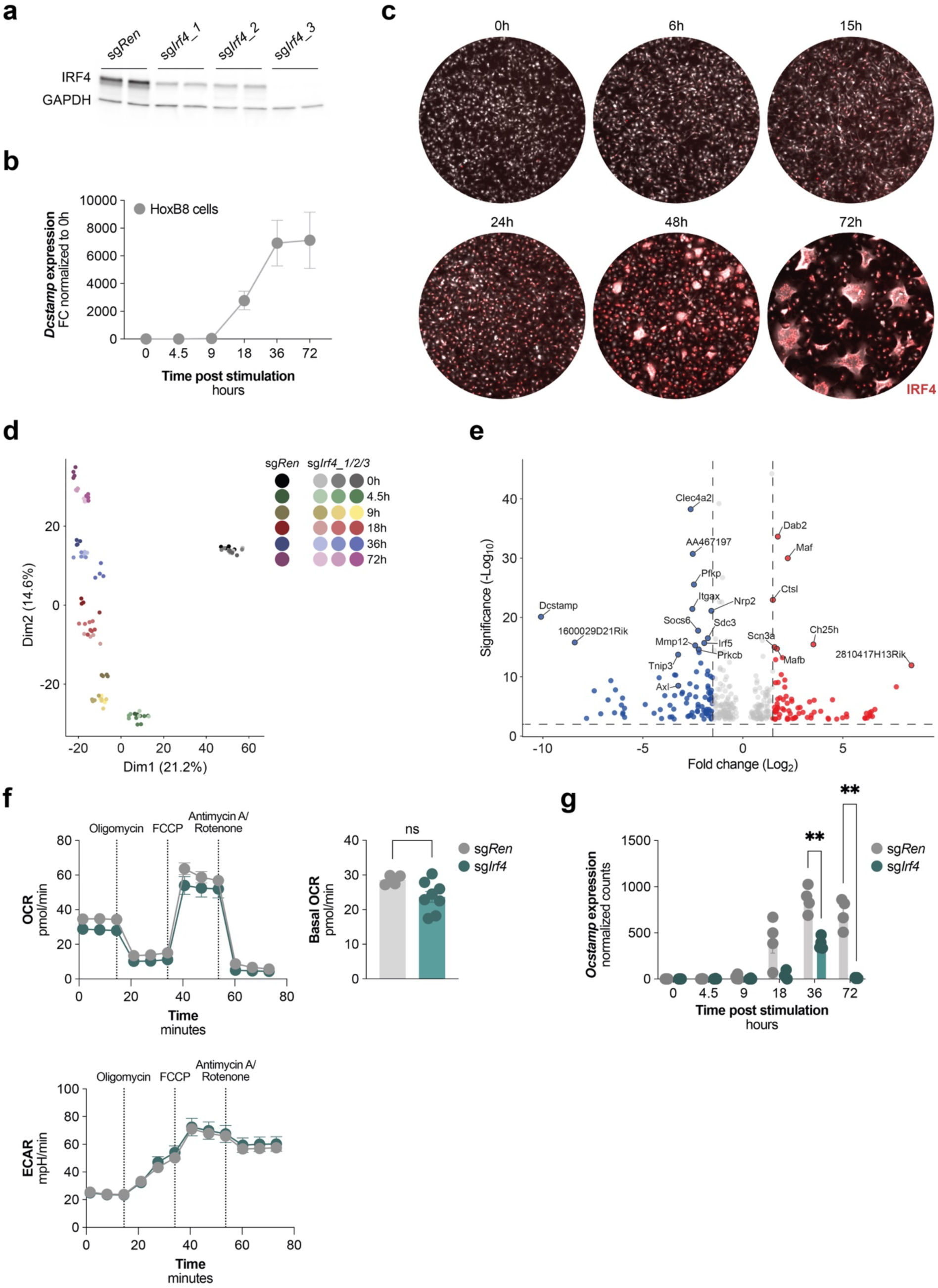
**(a)** Immunoblot of IRF4 and GAPDH (loading control) from protein lysates of *sgRen* and sg*Irf4_1/2/3* HoxB8 cells treated with IL-4 and GM-CSF for 3 days. **(b)** Expression levels of the MGC marker gene *Dcstamp* of sg*Ren* HoxB8 cells at various timepoints post fusion induction determined by qPCR. **(c)** Representative pictures of IRF4 stainings (AF555, red) of HoxB8 cells (anti-GFP, AF647, white) at different timepoints during MGC formation. **(d)** PCA plot of all bulk RNA-seq samples generated from *sgRen* and sg*Irf4_1/2/3* HoxB8 at different timepoints post fusion induction. **(e)** Volcano plot depicting DEGs of sg*Ren* vs sg*Irf4_1* at the 18 hours timepoint. Significantly up- and downregulated genes based on adjusted p-value of 0.05 are shown in red and blue, respectively. **(f)** Oxygen consumption rate (OCR) and extracellular acidification rate (ECAR) of MGC precursor cells derived from either *sgRen* or sg*Irf4_1/3* HoxB8 cells 24h post fusion induction. **(g)** *Ocstamp* expression levels from bulk RNA-seq dataset of sg*Ren* and sg*Irf4_1* HoxB8 cells at different timepoints post fusion induction. **(b)** Data represent n=4 technical replicates per timepoint and are representative of 2 independent experiments. **(c)** Representative images of 2 individual experiments. **(d)** Data represent n=4 technical replicates per genotype (sg*Ren*, sg*Irf4_1/2/3*). **(e)** Data represent n=4 technical replicates per genotype, but only data from sg*Ren* and sg*Irf4_1* HoxB8 cells are shown. **(f)** Data represent n=4-8 technical replicates. Data of sgI*rf4_1* and sg*Irf4_3* have been pooled. **(g)** Data represent n=4 technical replicates per genotype, but only data from sg*Ren* and sg*Irf4_1* HoxB8 cells are shown. Data are mean ± SEM, **p < 0.01 two-way ANOVA post-hoc pairwise comparisons with Bonferroni correction.

## Suppl. Movies

**(1-4)** Movies guiding through thick lung granuloma sections of different fate mapping mouse strains (yellow, endogenous tdTomato signal; blue, Hoechst staining). Representatives of several granuloma images of in total n=3 biological replicates per genotype.

**(5-6)** Movies depicting the uptake of pHrodo (red, when phagocytosed) labelled apoptotic neutrophils and large neutrophil cell clumps of *in vitro* generated MGCs. Representatives of 5-9 images per well of n=2-3 technical replicates and 2 independent experiments.

